# Mutational Biases and Selection in Mitochondrial Genomes: Insights from a Comparative Analysis of Natural and Experimental Populations of *Caenorhabditis elegans*

**DOI:** 10.1101/2025.09.03.674070

**Authors:** Alexandre Schifano, Ulfar Bergthorsson, Vaishali Katju

## Abstract

Spontaneous mutations display biases in their relative frequencies with important consequences for genome structure and composition. While laboratory studies have provided important insights into the spontaneous mutation spectrum, laboratory environments for optimal growth may engender biases that are not representative of natural populations. We analyzed the mitochondrial genomes of 1,524 *Caenorhabditis elegans* natural isolates comprising 550 unique haplotypes to investigate mtDNA polymorphism in the wild. Ancestral reconstruction was used to polarize 2,464 variants (88 indels, 2,376 SNPs) and the results were compared to mutations identified in experimental lines under relaxed selection. MtDNA variant distribution in natural isolates is strongly dependent on site-degeneracy in a manner consistent with purifying selection. There is significant variation in the synonymous and nonsynonymous polymorphism between genes. Specifically, ETC complex I genes are enriched for nonsynonymous polymorphism. The probability of synonymous mutation is higher at sites with flanking G/C nucleotides and the per gene synonymous polymorphism is negatively correlated with A+T-content at the 1^st^ and 2^nd^ codon positions. Furthermore, the 5′ and 3′ ends of genes have both higher A+T-content and less synonymous polymorphism than central regions. There is evidence of natural selection for preferred codons. We identify the first cases of large heteroplasmic mtDNA structural variants in *C. elegans* natural isolates, comprising deletions and duplications. Although some patterns of mtDNA mutational bias are similar between laboratory and natural populations, there exist significant differences. In particular, transversions typically associated with oxidative damage are less common at four-fold degenerate sites in natural populations relative to the laboratory.

**Significance:** Elements of genome structure, such as base composition and DNA strand bias are in large parts caused by the spectrum of spontaneous mutations. However, laboratory estimates of the mutation spectrum can be biased in an artificial environment that favors fast growth and short generation times. Polarized synonymous and rare variants in the mitochondrial genome of 1,524 natural isolates of *Caenorhabditis elegans* were analyzed to test if laboratory estimates of the mutation spectrum are representative of spontaneous mutations in the wild. The genomic distribution of synonymous variants is influenced by (i) base composition differences between and within genes, and (ii) natural selection for preferred codons. Certain categories of mutations, especially G/C → T/A transversions, are underrepresented in natural isolates relative to laboratory experiments. This discrepancy suggests that oxidative DNA damage is not as important a source of mutations in the wild relative to laboratory populations.

## Introduction

The acquisition of mitochondria was a crucial stage in eukaryotic evolution, facilitating gains in energy production through oxidative phosphorylation (Lane 2011). Most animal mitochondria are maternally inherited and comprise small compact genomes with higher mutation rates than nuclear DNA (Brown et al. 1979; Ladoukakis and Zouros 2017). Furthermore, animal mtDNA seem to undergo very little to no recombination (Eyre-Walker et al. 1999). Hence, their evolution is essentially clonal and new mitochondrial genotypes (mitotypes) are created by the accumulation of mutations and not by recombination. The study of mtDNA mutation spectra provide insights into the influence of spontaneous mutations on mtDNA genome evolution by helping decipher the origin of observed DNA base composition and strand biases, as well as the respective roles of both natural selection and mutational biases in shaping genomes, genome structure and codon usage. Furthermore, the base composition itself, a consequence of prior mutational biases and selection, can influence mutation rates. Finally, the mutation spectra can enable inferences about possible sources of mtDNA mutations and link them to both intrinsic and environmental factors promoting such mutations. For example, the consequences of reactive oxygen species (ROS) are predicted to have specific mutagenic effects which might be especially relevant when studying mtDNA mutations since ROS are by-products of mitochondrial activity (Kow 2002).

Despite a plethora of mutation accumulation (MA henceforth) which serve to minimize the influence of natural selection on newly originating variants, only a small subset have analyzed the rate and spectrum of spontaneous mutations of mitochondrial origin (Denver et al. 2000; Haag-Liautard et al. 2008; Lynch et al. 2008; Howe et al. 2009; Molnar et al. 2011; Saxer et al. 2012; Sung et al. 2012; Xu et al. 2012; Konrad et al. 2017; reviewed in Katju and Bergthorsson 2019). Because eukaryotic cells can host up to thousands of mtDNA copies, spontaneous mtDNA mutations will therefore initially only be present at a very low intracellular frequency compared to novel nuclear mutations. This difference makes the detection of novel mtDNA variants particularly challenging with the use of common DNA sequencing methods and necessitating extensive MA experiments over multiple generations to allow new mtDNA variants to increase to detectable frequencies. The fate of most spontaneous mutations is loss by genetic drift. Hence, even in MA experiments maintained across multiple parallel lines for multiple generation, the number of detectable mutations remains low with regular sequencing methods (Sanger or Illumina HiSeq), compromising their statistical power to analyze the mtDNA mutation spectrum. Furthermore, the diminutive size of animal mtDNA genomes exacerbates this challenge of detecting novel mtDNA genomes in MA experiments even though animal mtDNA mutation rates can be several orders of magnitude greater than their nuclear counterparts (Konrad et al. 2017).

Several studies have attempted to measure the mtDNA mutation spectrum in the model nematode, *Caenorhabditis elegans*, often employing an MA approach in order to minimize the influence of natural selection (Denver et al. 2000; Konrad et al. 2017; Waneka et al. 2021; Leuthner et al. 2022), but the results have varied substantially between studies. Highly sensitive sequencing methods such as duplex sequencing enable the detection of even the rarest variants with high fidelity (Schmitt et al. 2012) and were employed by two recent studies focusing on *C. elegans* mtDNA mutations (Waneka et al. 2021; Leuthner et al. 2022). Although duplex sequencing allowed for a far greater yield of spontaneous mtDNA mutations in these two recent studies, their results differed somewhat despite the employment of the same sequencing technology. The results from duplex sequencing reported by Waneka et al. (2021) are consistent with the MA experiment in Konrad et al. (2017), with both studies detecting a strong G/C → A/T mutational bias, with G/C → A/T transitions being the most common, followed by G/C → T/A transversions. Waneka et al. (2021) additionally observed a strand bias for suspected oxidative damage on the exposed strand. In contrast, a duplex sequencing study by Leuthner et al. (2022) found that mtDNA mutations increasing A+T-content were predominantly transversions and G/C → C/G mutations, which were rare or absent in preceding studies, were the second most common class of base substitutions. The large number of mtDNA mutations detected in both duplex sequencing studies argues against chance events as being the source of the differences between them. Konrad et al. (2017) additionally analyzed the mtDNA genomes of 38 *C. elegans* wild isolates (Thompson et al. 2013) to investigate the spectrum of mtDNA mutations observed under selective conditions in comparison to spontaneous mtDNA mutations originating in their MA experiment under a regime of minimal selection.

The signature of oxidative damage across several laboratory-based *C. elegans* studies may appear to support the hypothesis that mitochondrial ROS are a major contributor to mtDNA mutations. Indeed, one hypothesis regarding oxidative damage posits that higher metabolic demands leading to increase ATP production may contribute to the production of ROS (Gillooly et al. 2005) and therefore increase G/C → T/A transversions. Can optimal growth conditions for *C. elegans* in the laboratory be a factor in promoting oxidative damage relative to wild populations? Our understanding of the mtDNA mutation rate in *C. elegans* is primarily based on laboratory experiments with worms cultivated at the maximum rate in a predictable environment (Denver et al. 2000; Konrad et al. 2017; Waneka et al. 2021; Leuthner et al. 2022). In contrast, wild worms can have vastly different growth rates and generation times, and experience a diversity of growth conditions and environmental stressors not encountered under laboratory conditions. These in turn may influence DNA replication and repair resulting in mutation patterns that differ from laboratory evolved lines. One of the main objectives of this study is to test if the spectrum of mtDNA mutations in natural populations of *C. elegans* differs from that of laboratory studies. To this end, we examined the mtDNA genomic sequences of 1,524 natural isolates of *C. elegans* from the CaeNDR database (Crombie et al. 2024) representing 550 genetically distinct isotypes clustered into 347 unique mitotypes. We employed a novel programming algorithm for high accuracy ancestral sequence reconstruction (ASR) for base substitutions and small indel changes (Jowkar et al. 2022) in order to identify and polarize mtDNA variants occurring in these *C. elegans* natural isolates. Natural populations are subject to stronger purifying selection than laboratory populations, but with a large data set which includes 900 mutations at four-fold degenerate sites, we demonstrate significant differences in the mutation spectrum between natural and laboratory populations. In addition to conducting tests for codon usage bias, neighbor effect, and positive selection, we additionally identify the first instances of several large mitochondrial deletions and duplications in natural populations of *C. elegans*.

## Results

We identified and polarized mtDNA mutations from 531 distinct mtDNA isotypes (550 minus the 19 strains flagged for possible mtDNA contamination) corresponding to 1,524 natural isolates of *C. elegans* (supplementary file S1) using a method combining phylogeny and ancestral sequence reconstruction. With the exception of two low-complexity regions comprising 466 bp in the reference mitochondrial genome, the entire mitochondrial genome (13,315 bp in the Bristol N2 laboratory strain) was analysed. We identified 2,464 mutations comprising 88 small indels and 2,376 SNPs (supplementary file S2). Approximately 46% (1,125) of these mutations were inferred between nodes within the phylogeny and the remaining 54% (1,339) were identified in the terminal branches when compared to their most recent reconstructed ancestor. The number of mutations per strain or node varied from 0 to 101 with a mean and median count of five and two mutations per branch, respectively. The largest number of inferred mutations along a branch was located in the ancestor of the Hawaiian cluster which includes the natural isolate CB4856. Transitions accounted for 81.8% of all SNPs, which included 1,062 A/G → G/C and 881 G/C → A/G mutations (Table 1). The transition to transversion (*Ts/Tv*) ratio for all SNPs is 4.49. The most common transversion was A/T → T/A (226) followed by G/C → T/A (144), A/T → C/G (54) and G/C → C/G (9).

**Table 1.** The number and normalized proportions of base substitutions in this study and comparisons to three preceding experimental studies in a laboratory setting. The normalized proportions of each mutation were computed by first dividing the counts of a given mutation type (for example, A/T → G/C) by the number of the mutated nucleotides (A/T) in the genome. The results were then divided by the sum of these fractions for all six mutational categories.

| Study | Mutation | Transversions |  |  |  | Transitions |  |
| --- | --- | --- | --- | --- | --- | --- | --- |
| | | A/T → C/G | A/T → T/A | G/C → T/A | G/C → C/G | A/T → G/C | G/C → A/T |
| This study | Counts | 54 | 224 | 144 | 9 | 1057 | 875 |
|  | Proportions | 0.023 | 0.095 | 0.061 | 0.004 | 0.447 | 0.370 |
|  | Normalised proportions | 0.012 | 0.050 | 0.098 | 0.006 | 0.233 | 0.601 |
| This study<br>Four-fold degenerate sites | Counts | 39 | 129 | 48 | 5 | 385 | 290 |
|  | Proportions | 0.044 | 0.144 | 0.054 | 0.006 | 0.430 | 0.324 |
|  | Normalised proportions | 0.014 | 0.047 | 0.111 | 0.012 | 0.141 | 0.675 |
| Waneka *et al.* (2021) | Counts | 2 | 20 | 79 | 8 | 50 | 94 |
|  | Proportions | 0.008 | 0.079 | 0.312 | 0.032 | 0.198 | 0.372 |
|  | Normalised proportions | 1.25E-05 | 0.032 | 0.388 | 0.039 | 0.079 | 0.462 |
| Waneka *et al.* (2021)<br>Four-fold degenerate sites | Counts | 1 | 5 | 4 | 0 | 10 | 9 |
|  | Proportions | 0.034 | 0.172 | 0.138 | 0 | 0.345 | 0.310 |
|  | Normalised proportions | 0.010 | 0.051 | 0.258 | 0 | 0.101 | 0.580 |
| Leuthner *et al.* (2022) | Counts | 22 | 76 | 300 | 225 | 44 | 93 |
|  | Proportions | 0.029 | 0.100 | 0.395 | 0.296 | 0.058 | 0.122 |
|  | Normalised proportions | 1.42E-05 | 0.037 | 0.457 | 0.343 | 0.022 | 0.142 |
| Konrad *et al.* (2017) | Counts | 0 | 1 | 3 | 0 | 0 | 5 |
|  | Proportions | 0 | 0.111 | 0.333 | 0 | 0 | 0.556 |
|  | Normalised proportions | 0 | 0.039 | 0.360 | 0 | 0 | 0.600 |

### Mutational spectra of mtDNA base substitutions in natural isolates differ from that of experimental lines

We inferred the mutation spectra using all the mtDNA base substitutions found in this study (Fig. 1a) as well as one using only base substitutions found at four-fold degenerate sites (Fig. 1b). We also adapted the data from two recent studies investigating *C. elegans* mitochondrial mutations under experimental laboratory conditions employing the wildtype Bristol (N2) strain (Waneka et al. 2021; Leuthner et al. 2022) (Fig. 1c and d). The number of mutations was normalized by the base composition of the reference genome (or of the reference genome’s four-fold degenerate sites). In other words, the counts of G/C → N/N mutations were divided by the number of G/C base pairs in the reference genome (and all A/T → N/N mutations by the number of A/T base pairs) to account for the inherent A/T-bias in the *C. elegans* mitochondrial genome.

**Fig. 1.**
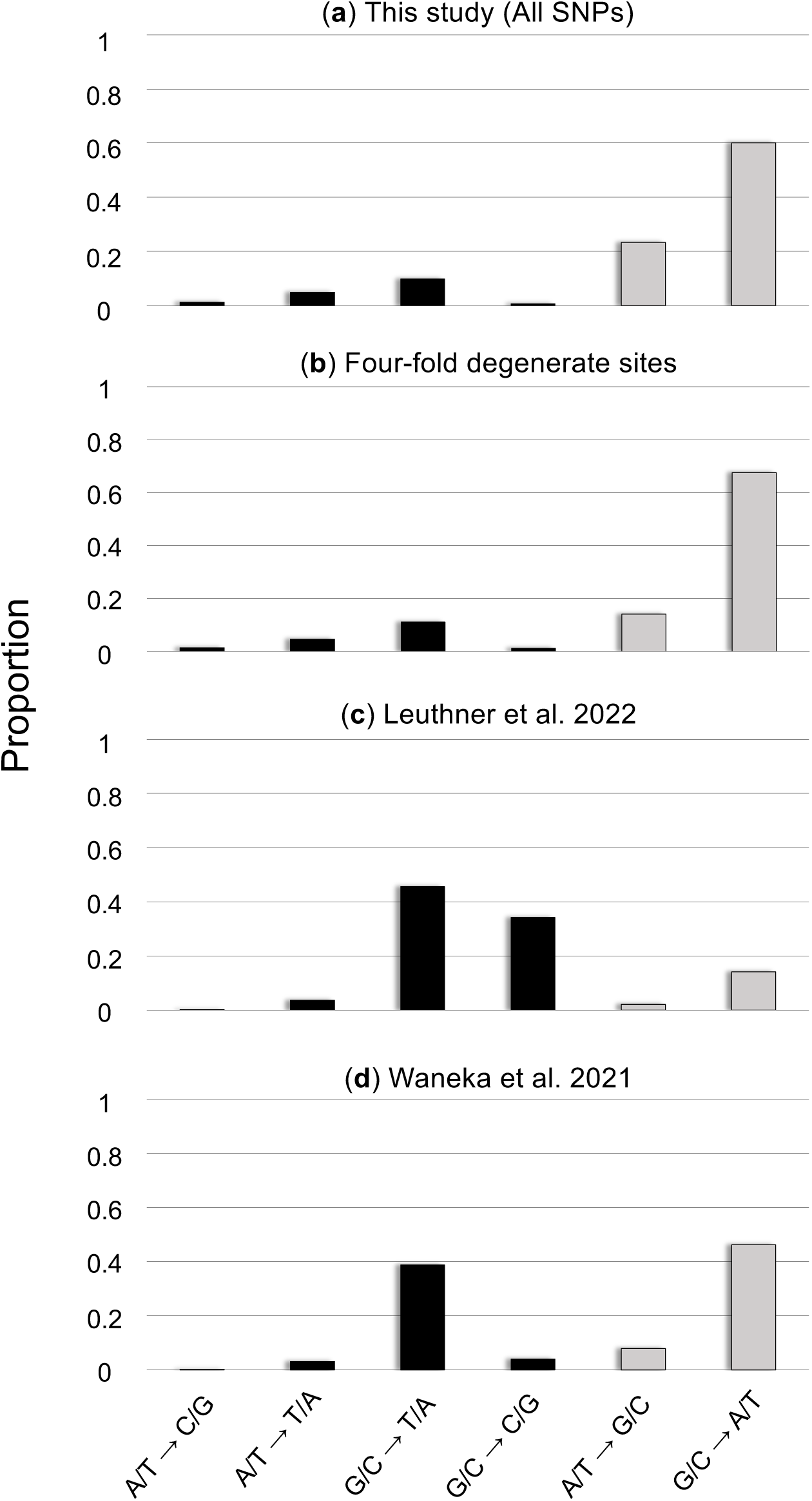
The mutational spectrum of base substitutions in the mitochondrial genome of *C. elegans*. The mutation counts were normalized by the reference genome’s base composition and scaled such that the sum of the frequencies in each experiment equals one. Transitions and transversions are denoted in grey and black, respectively. a) All 2,376 base substitutions inferred in this study of natural isolates. b) Base substitutions at four-fold degenerate sites in the natural isolates (900 SNPs). c) Base substitutions reported in a duplex sequencing study by Leuthner et al. 2022 (760 SNPs). d) Base substitutions reported in a duplex sequencing study by Waneka et al. 2021 (253 SNPs).

A key difference between natural isolates and the results from duplex sequencing of laboratory strains is that transitions are significantly more frequent in the wild compared to the laboratory experiments (Table 1; Fig. 1a). A comparison of the number of transitions and transversions between four-fold degenerate sites in natural isolates (Fig. 1b) and variants at all sites in duplex sequencing from laboratory experiments (Waneka et al. 2021; Leuthner et al. 2022; Fig. 1c and d) is highly significant (χ*^2^* test-statistic = 548.68; *d.f.* = 2; *N* = 1,913; *p* = 7.2 × 10^-120^). The results from Leuthner et al. (2022) are clearly an outlier here with only 18% of all mutations identified as transitions (Fig. 1c). If we only compare four-fold degenerate sites in natural isolates (Fig. 1b) and the duplex sequencing variants at all sites of Waneka et al. (2021) (Fig. 1d) where transitions comprised 75% and 57% of all mutations, respectively, the difference is still highly significant (χ*^2^* test-statistic = 33.18; *d.f.* = 1; *N* = 1,153; *p* = 8.4 × 10^-9^).

G/C → A/T transitions are more frequent than A/T → G/C at all sites in duplex sequencing experiments whereas the converse is observed for four-fold degenerate sites in the natural isolates (Table 1; χ*^2^* test-statistic = 44.26; *d.f.* = 2; *N* = 960; *p* = 2.5 × 10^-10^). However, these transitions still show an AT-bias as the ratio of 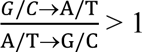 when the base composition of the genome is accounted for in both the laboratory experiments and natural isolates (Table 1). The ratio between these two transitions is higher in the duplex sequencing experiments where the G/C → A/T transitions per G/C base pair are 5.8 − 6.5× more likely than A/T → G/C transitions per A/T base pair. In contrast, the G/C → A/T transitions are only 2.6× more likely than A/T → G/C transitions when controlled for base composition (Table 1) in the natural isolates. The 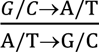 ratio in the natural isolates is observed to be 4.8 at four-fold degenerate sites which demonstrates that the high number of A/T → G/C transitions at these sites is due to the high A+T-content at four-fold degenerate sites of the mitochondrial genome (Table 1).

Furthermore, there are also significant differences between the natural isolates and the duplex sequencing of laboratory strains in the relative numbers of different transversions (Table 1; χ*^2^* test-statistic = 338.82; *d.f.* = 6; *N* = 953; *p* = 3.9 × 10^-70^). To investigate differences in the relative proportions of the various transversion classes, our analysis was restricted to four-fold degenerate sites from natural isolates as transversions at two-fold degenerate sites of protein-coding sequences result in amino acid replacement which are subject to stronger purifying selection pressure in the wild. Similar to that observed previously in the case of transitions, one of the duplex sequencing studies (Leuthner et al. 2022) is a strong outlier here as it reports a high proportion of G/C → C/G transversions, a substitution class that is usually amongst the rarest substitutions in all other studies of mtDNA mutation rates in *C. elegans*, and ranging between 0% to 4% of all mutations. If we restrict the comparison to four-fold degenerate sites in the natural isolates and the duplex sequencing experiments of Waneka et al. (2021), the difference in the proportions of various types of transversion remains highly significant (Table 1; χ*^2^* test-statistic = 94.23; *d.f.* = 3; *N* = 330; *p* = 2.7 × 10^-20^). Although the four categories of transversions listed in Table 1 are in strikingly different proportions between four-fold degenerate sites in natural isolates and the duplex sequencing study of Waneka et al. (2021), we note that the most common transversions in the laboratory study are G/C → T/A (72.5% of all transversions, 31.2% of all base substitutions) which were attributed to oxidative damage. In contrast, the very same G/C → T/A transversions are only 21.7% and 5.3% of all transversions and base substitutions at four-fold degenerate sites in the natural isolates, respectively.

### Base composition of four-fold degenerate sites in natural isolates is not in mutational equilibrium

At mutational equilibrium, the number of mutations that increase the A+T-content of the mitochondrial genome is expected to be equal to the number of G/C-increasing mutations. In both of the duplex sequencing studies (Waneka et al. 2021; Leuthner et al. 2022), A/T-increasing mutations vastly outnumber G/C-increasing mutations (Table 1). The ratios of A/T-increasing to G/C-increasing mutations were 6.0 and 3.3 in the studies by Leuthner et al. (2022) and Waneka et al. (2021), respectively. However, the base composition at nondegenerate sites is not expected to be in mutational equilibrium due to selection on the amino acid sequence. Consequently, nondegenerate sites in protein-coding regions of mtDNA typically have higher G+C-content than two-fold and four-fold degenerate sites. The high ratio of A/T-increasing to G/C-increasing mutations under relaxed selection in laboratory experiments is a reflection of the deviation from mutational equilibrium at nondegenerate sites. For instance, if we only consider the four-fold degenerate sites in Waneka et al. (2021), the ratio of A/T-increasing mutations to G/C-increasing mutations is closer to one (1.18). In the natural isolates, the numbers of A/T-increasing and G/C-increasing mutations are more similar, with a ratio of 0.92. However, the ratio of A/T-increasing to G/C-increasing mutations is only 0.80 at four-fold degenerate sites in the natural isolates. The difference between the number of A/T-increasing and G/C-increasing mutations at four-fold degenerate sites in natural isolates is significant (χ*^2^* test-statistic = 9.66; *d.f.* = 1; *N* = 766; *p* = 0.0019). Furthermore, there is no significant difference between the ratio of A/T-increasing to G/C-increasing mutations inferred at the four-fold degenerate sites within the internal nodes of the phylogenetic tree (0.84; *N* = 406) versus single mitotypes (0.76; *N* =360) within the natural isolates (χ*^2^* test-statistic = 0.48; *d.f.* = 1; *N* = 766; *p* = 0.4851). The observation that G/C-increasing mutations significantly outnumber A/T-increasing mutations at four-fold degenerate sites in the wild suggests that these sites are not in mutational equilibrium.

### Strong signature of purifying selection in mtDNA protein-coding genes of natural isolates

Polymorphic sites with respect to base substitutions in the mtDNA protein-coding regions of the natural isolates show evidence of strong purifying selection acting on the first and second codon positions (Fig. 2a). The second codon position, which is always nondegenerate, has only ∼7% of the total number of observed mtDNA base substitutions. The first and third codon positions harbor approximately 19% and 74% of base substitutions in the mtDNA protein-coding genes, respectively. In the duplex sequencing study by Waneka et al. (2021), the proportions of mutations per codon position are closer to the neutral expectation under a regime of relaxed selection (Fig. 2a). The slightly higher proportions of mutations at the first and second codon positions in their study possibly reflects higher G/C-content at these sites rendering them more prone to mutations. With regards to codon degeneracy, we standardized the observed number of base substitutions in mtDNA protein-coding regions by the total number of nondegenerate, two-fold degenerate and four-fold degenerate sites in the *C. elegans* reference mtDNA genome. The relative proportion of base substitutions observed in the natural isolates at four-fold degenerate sites was 0.64, compared with 0.05 and 0.31 at nondegenerate and two-fold degenerate sites, respectively (Fig. 2b). These contrasts are even more evident when comparing the natural isolates’ data with the results from Waneka et al. (2021), where four-fold and nondegenerate sites harbor comparable proportions of mutations (Fig. 2b). Interestingly, the results adapted from the study by Waneka et al. (2021) find two-fold degenerate sites exhibiting a reduction in the proportion of mutations compared to four-fold and nondegenerate sites (Fig. 2b).

**Fig. 2.**
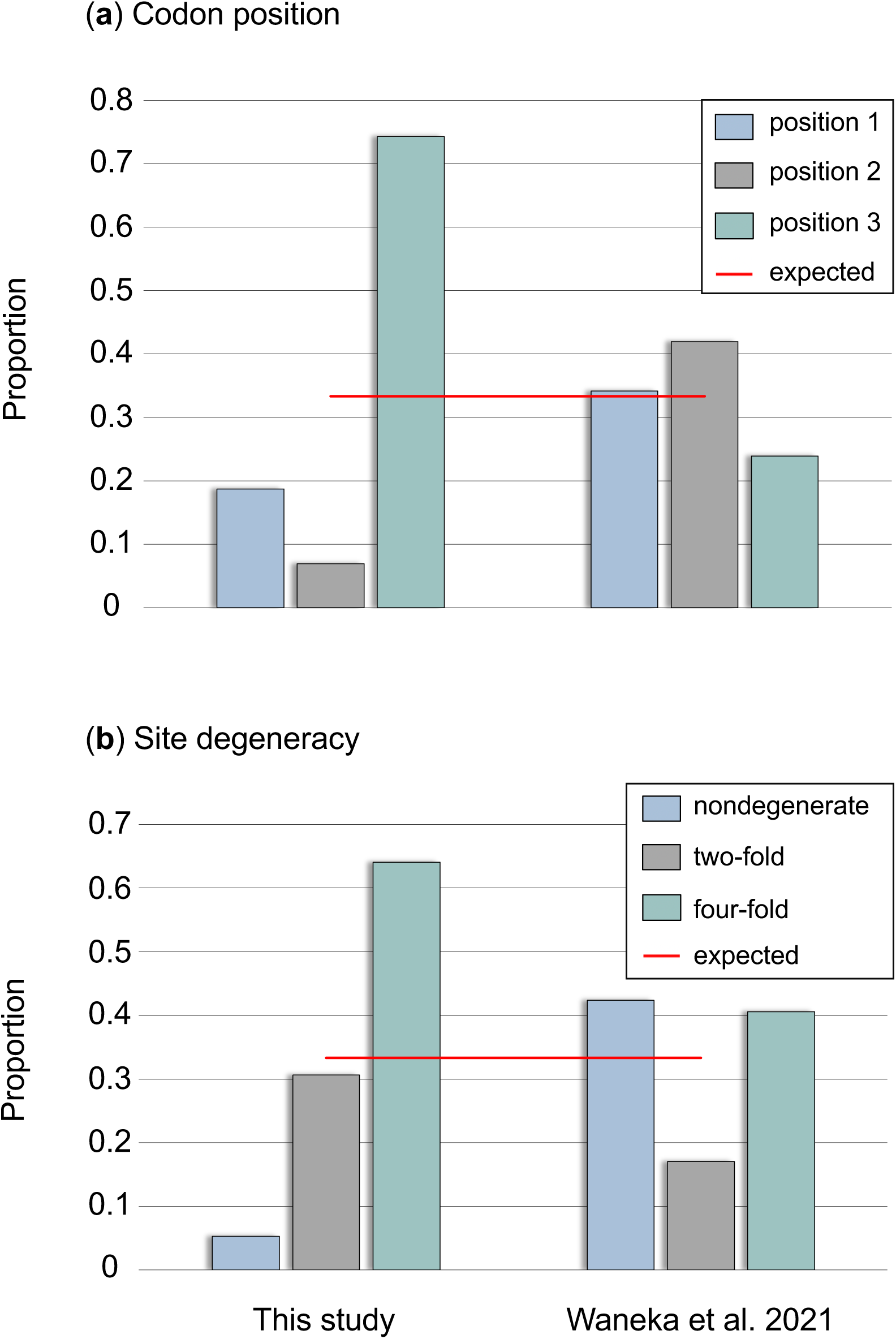
The proportion of polymorphic base substitutions categorized by codon position and site degeneracy. a) The proportion of polymorphic base substitutions at the first, second and third codon position in *C. elegans* natural isolates (left) and in duplex sequencing results under relaxed selection (Waneka et al. 2021, right). b) The proportion of polymorphic base substitutions at nondegenerate, two-fold and four-fold degenerate sites in the *C. elegans* natural isolates were standardized by the total number of nondegenerate, two-fold and four-fold degenerate sites in the *C. elegans* mtDNA reference genome (left) and duplex sequencing results under relaxed selection (Waneka et al. 2021, right). The red horizontal lines represent the expected proportions if all sites accumulate base substitutions at equal rates.

The *Ts/Tv* ratios of inferred base substitutions vary predictably between codon positions in the natural isolates with the third and second codon position displaying the highest (5.35) and lowest ratio (2.66), respectively (Fig. 3a). For comparison, in the duplex sequencing results of Waneka et al. (2021) the mutation proportions per codon position are much closer to the neutral expectation under relaxed selection (Fig. 3a). Overall, the *Ts/Tv* ratios per codon position are lower in the duplex sequencing study (Waneka et al. 2021), demonstrating that the bias towards transitions is less pronounced under a regime of relaxed selection. Surprisingly, the *Ts/Tv* ratio at the third codon position in the Waneka et al. (2021) study is more than 2× higher than that observed at the first and second codon positions (Fig. 3a). In the natural isolates, transversions comprise ∼6% of the mutations at two-fold degenerate sites while constituting approximately 25% and 31% of mutations at four-fold degenerate and nondegenerate sites, respectively. The *Ts/Tv* ratio varies by site degeneracy in the protein-coding genes, with two-fold degenerate sites having the highest *Ts/Tv* ratio as expected (16.70), followed by four-fold degenerate sites (3.07) and nondegenerate sites (2.25) (Fig. 3b). Although the *Ts/Tv* ratios are far lower in the duplex sequencing study of rare new mutations under a regime of relaxed selection (Waneka et al. 2021), two-fold degenerate sites still display a higher *Ts/Tv* ratio than four-fold degenerate and nondegenerate sites (Fig. 3b), indicative of a modicum of purifying selection operating in this experimental regime.

**Fig. 3.**
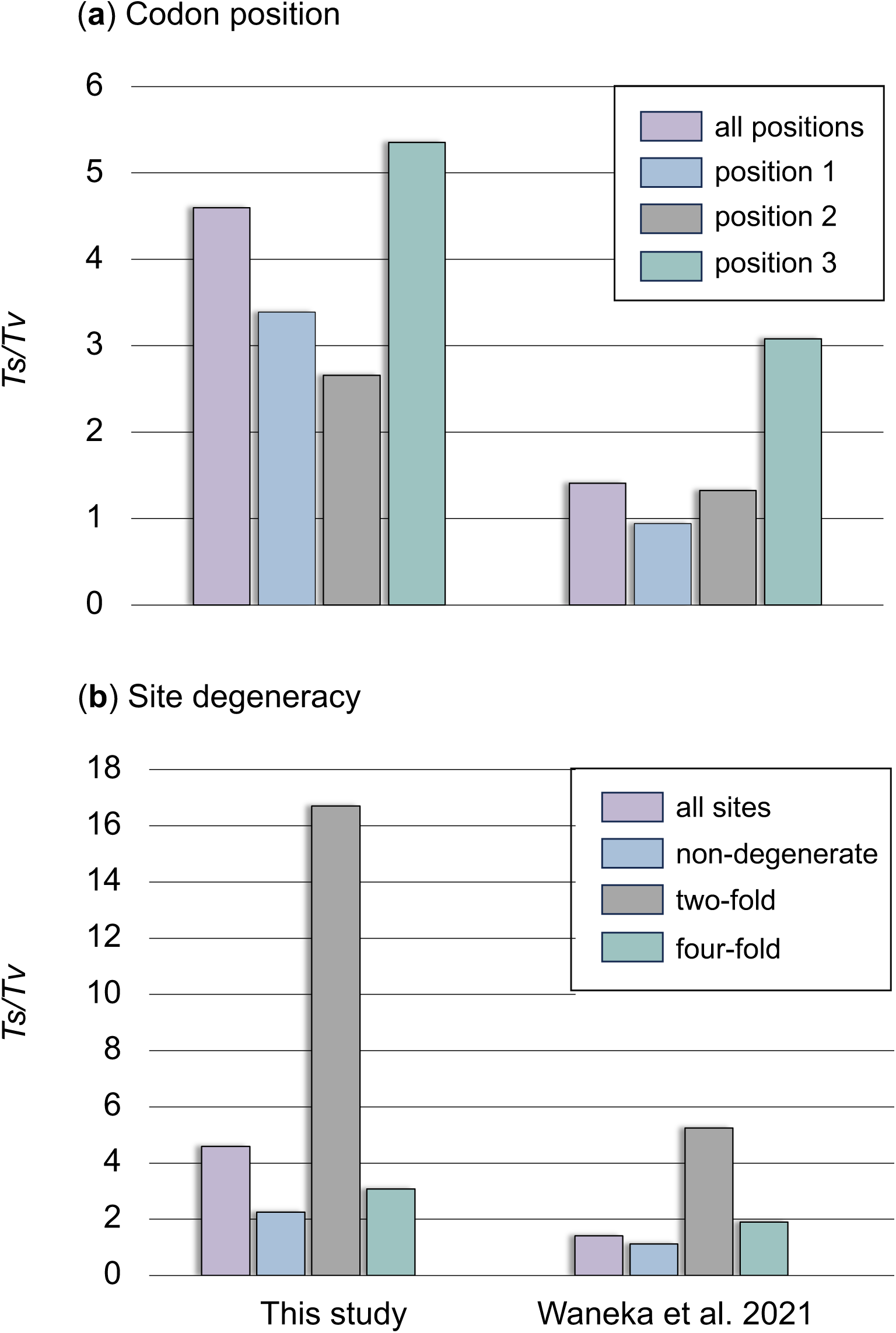
Transition to transversion (*Ts/Tv*) ratios of base substitutions by codon position and site degeneracy. a) The *Ts/Tv* ratios at different codon positions in *C. elegans* natural isolates (left) and in duplex sequencing results under relaxed selection (Waneka et al. 2021, right). b) The *Ts/Tv* ratios at all sites, non-, two-fold, and four-fold degenerate sites in *C. elegans* natural isolates (left) and in duplex sequencing results under relaxed selection (Waneka et al. 2021, right).

### Numbers of polymorphic base substitutions vary by mtDNA gene and ETC complex

Of the 12 protein-coding genes comprising the mitochondrial genome of *C. elegans*, *nd4L* has significantly fewer polymorphic substitutions (*G* test-statistic = 11.1, *d.f.* = 1, *p* = 8.41×10^-4^) whereas *nd1* has significantly more polymorphic substitutions (*G* test-statistic = 31.7, *d.f.* =1, *p* = 1.79×10^-8^) relative to the other protein-coding genes (Fig. 4a). Furthermore, the nonsynonymous to synonymous (*N*/*S*) ratios are significantly higher in *nd*2 (Fisher’s exact test: *p* = 3.44 × 10^-3^) and in *nd6* (Fisher’s exact test: *p* = 0.0351) but significantly lower in *cox1* (Fisher’s exact test: *p* = 4.06 × 10^-4^) compared to the other protein-coding genes (Fig. 4b). In *nd2*, there were both fewer synonymous mutations (*G* test-statistic = 9.95, *d.f.* = 1, *p* = 1.61 × 10^-4^, Fig. 4c) and more nonsynonymous mutations (*G* test-statistic = 11.14, *d.f.* = 1, *p* = 8.44 × 10^-4^, Fig. 4d) compared to other mtDNA protein-coding genes. Similarly, the low *N/S* in *cox1* is due to both fewer nonsynonymous mutations (*G* test-statistic = 18.87, *d.f.* = 1, *p* = 1.40 × 10^-5^, Fig. 4d) and more synonymous mutations than the genomic average. The high *N/S* in *nd6* is due to significantly greater number of nonsynonymous mutations compared to the genomic average (*G* test-statistic = 10.81, *d.f.=* 1, *p* = 1.01 × 10^-3^; Fig. 4d). The gene bearing the most mutations per site, *nd1* (Fig. 4a), has significantly greater number of mutations at both synonymous (*G* test-statistic = 15.54, *d.f.* = 1, p = 1.01 × 10^-5^) and nonsynonymous sites (*G* test-statistic = 21.09, *d.f.* = 1, *p* = 4.37 × 10^-6^; Figs. 4c and d).

**Fig. 4.**
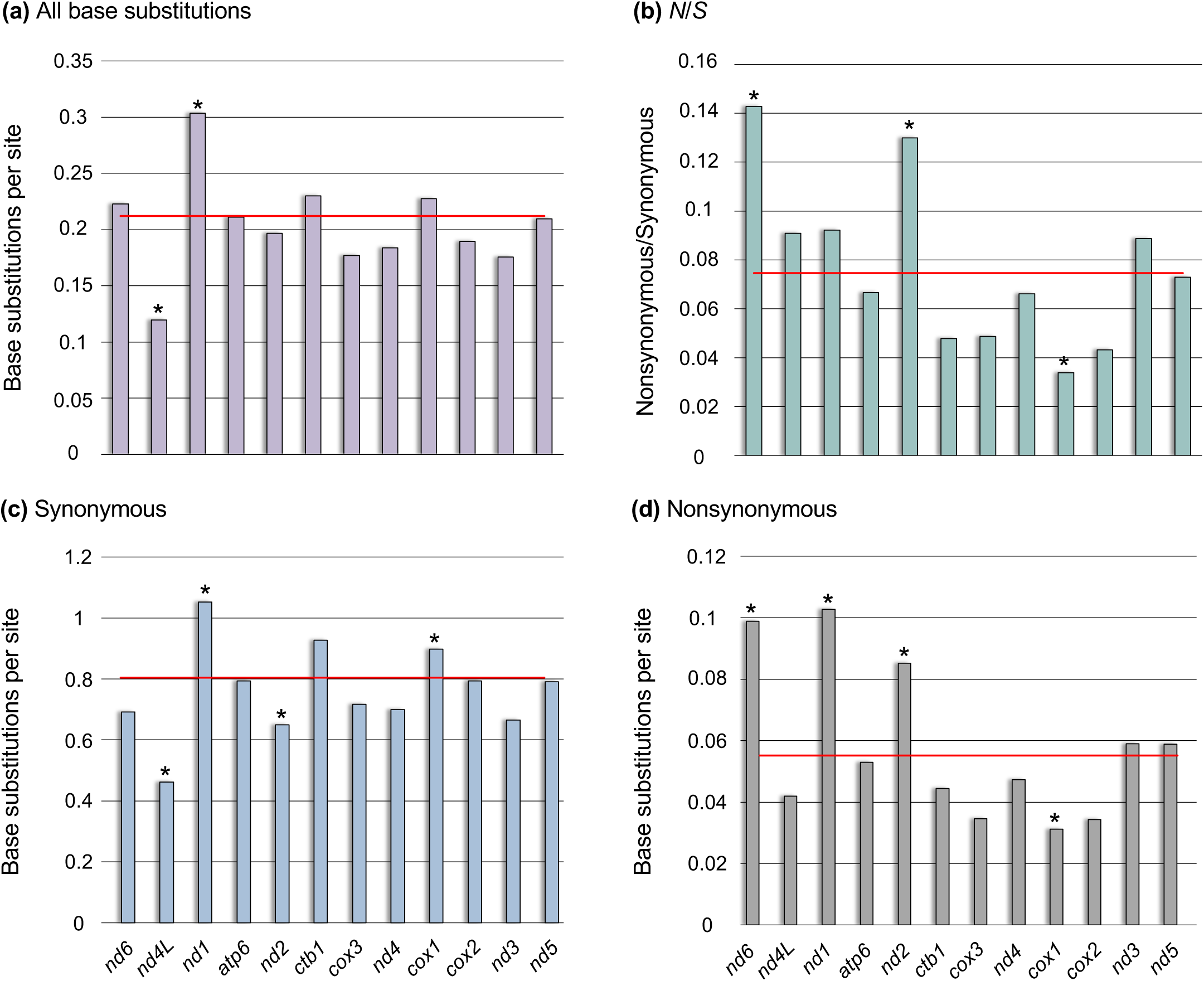
The frequency of polymorphic base substitutions per site in the 12 mtDNA protein-coding genes in the *C. elegans* natural isolates. a) The frequency of all polymorphic base substitutions. b) The nonsynonymous to synonymous ratio (*N*/*S*) per gene. c) The number of synonymous substitutions per synonymous site. d) The number of nonsynonymous substitutions per nonsynonymous site. The red horizontal lines represent the average for the protein-coding mitochondrial genome. *G*-tests comparing the observed versus expected mutation counts under homogeneous mutation frequency were performed using a Bonferroni correction for multiple comparisons (*p*-value < 0.004) except for the nonsynonymous to synonymous ratios which utilized Fisher’s exact tests. Significant deviations from the expected frequencies are denoted with an asterisk.

There were no significant differences in the total number of mutations per site when the 12 mtDNA protein-coding genes were grouped into their respective electron transport chain (ETC) complexes (*G* test-statistic = 2.37, *d.f.* = 1, *p* = 0.498; Fig. 5a). However, genes in complex I display a significantly higher *N/S* ratio (Fisher’s exact test: *p* = 1.93 × 10^-3^; Fig. 5b) due to both significantly fewer synonymous mutations (*G* test-statistic = 10.27, *d.f.* = 1, *p* = 1.35 × 10^-3^; Fig. 5c) and higher nonsynonymous mutations (*G* test-statistic = 36.28, *d.f.* = 1, *p* = 1.71 × 10^-9^; Fig. 5d) than genes in the other three mitochondrial respiratory complexes. Furthermore, the seven highest *N/S* ratios, even if not significantly departing from the genomic average, are found in ETC complex I genes. In contrast, mtDNA genes functioning in complex IV have a significantly lower *N/S* ratio (Fisher’s exact test: *p* = 1.40 × 10^-5^) owing to significantly fewer nonsynonymous mutations (*G* test-statistic = 33.93, *d.f.* = 1, *p* = 5.71 × 10^-9^; Fig. 5d) relative to the genes in other ETC complexes.

**Fig. 5.**
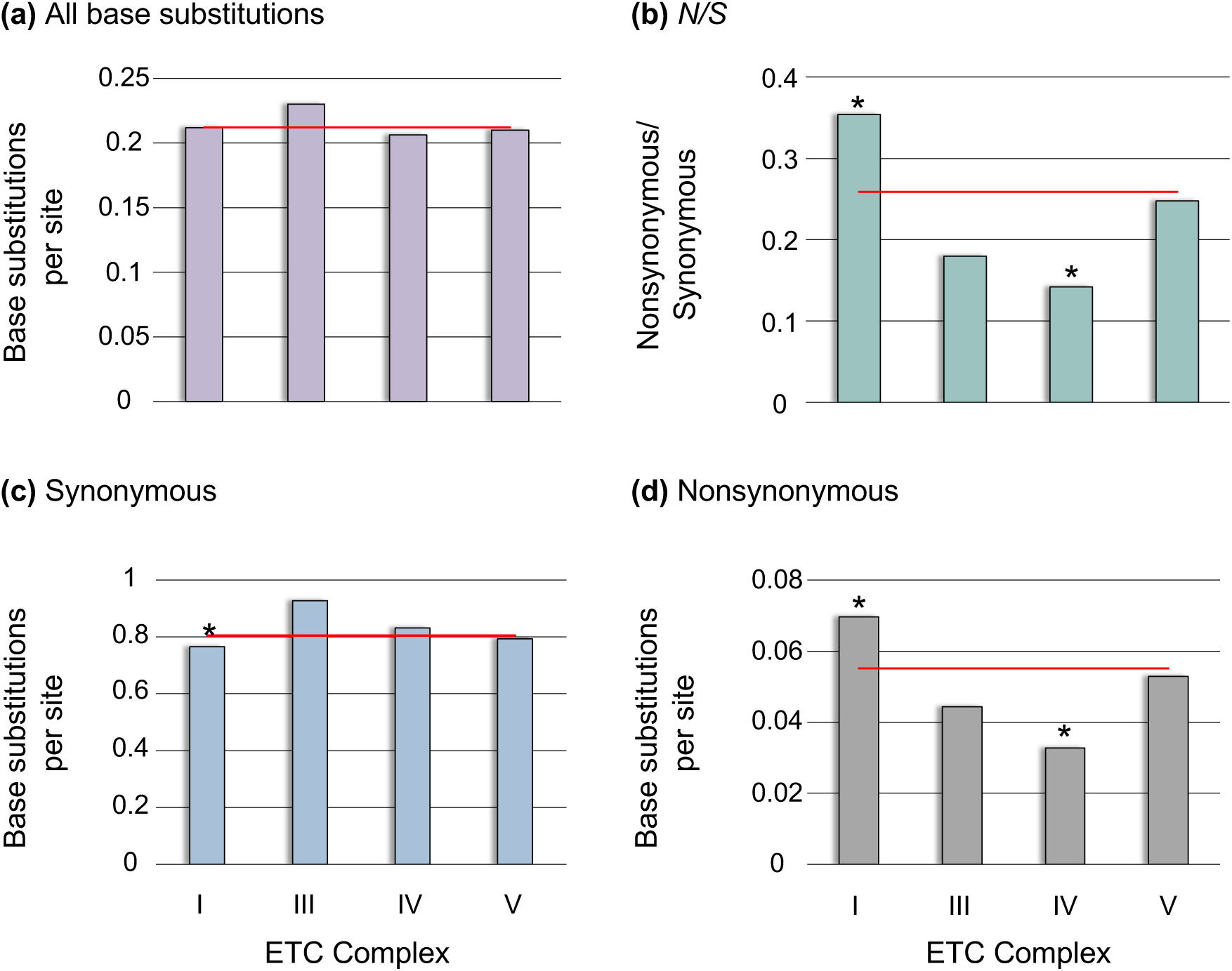
The frequency of polymorphic base substitutions per site with the mtDNA protein-coding genes grouped by their association with four electron transport chain (ETC) complexes. a) The frequency of all polymorphic base substitutions. b) The nonsynonymous to synonymous ratio (*N*/*S*) per ETC complex. c) The number of synonymous substitutions per synonymous site. d) The number of nonsynonymous substitutions per nonsynonymous site. The red horizontal lines represent the average for the protein-coding mitochondrial genome. *G*-tests comparing the observed versus expected mutation counts under homogeneous mutation frequency were performed using a Bonferroni correction for multiple comparisons (*p*-value < 0.0125) except for the nonsynonymous to synonymous ratios which utilized Fisher’s exact tests. Significant deviations from the expected frequencies are denoted with an asterisk.

### Local context-dependence of mutations

Sites with G/C flanking nucleotides, either as 5′ or 3′ neighbors, have higher probability of mutations than sites with A/T neighbors (Fig. 6). Significant departures from equal mutation distribution at two- and four-fold degenerate sites further support this observation (supplementary table S1). Two-fold degenerate sites have a 1.85× greater probability of a mutation if their 5′ neighbor is G/C compared to A/T. Likewise, there is a 1.58× greater probability of mutations at two-fold degenerate sites with 3′ G/C neighbors rather than A/T. Four-fold degenerate sites with 5′ G/C neighbors have a 1.7× greater probability of mutations relative to sites with 5′ A/T neighbors. Lastly, four-fold degenerate sites with 3′ G/C neighbors have a 1.91× greater probability of mutations compared to sites with 3′ A/T. Context-dependent mutations can also contribute to the variation in synonymous mutation rates between mtDNA genes. Because degenerate sites at 3^rd^ base position are flanked by nucleotides at 1^st^ and 2^nd^ codon positions, we tested for an association between base composition at these positions with synonymous polymorphisms. The synonymous polymorphisms in *C. elegans* mtDNA genes is positively correlated with the G+C-content at the 1^st^ and 2^nd^ codon positions (Pearson’s *r* = 0.703, *d.f.* = 10, *p* = 0.0107) but not at the 3^rd^ codon position (Pearson’s *r* = -0.002, *d.f.* = 10, *p* = 0.9958).

**Fig. 6.**
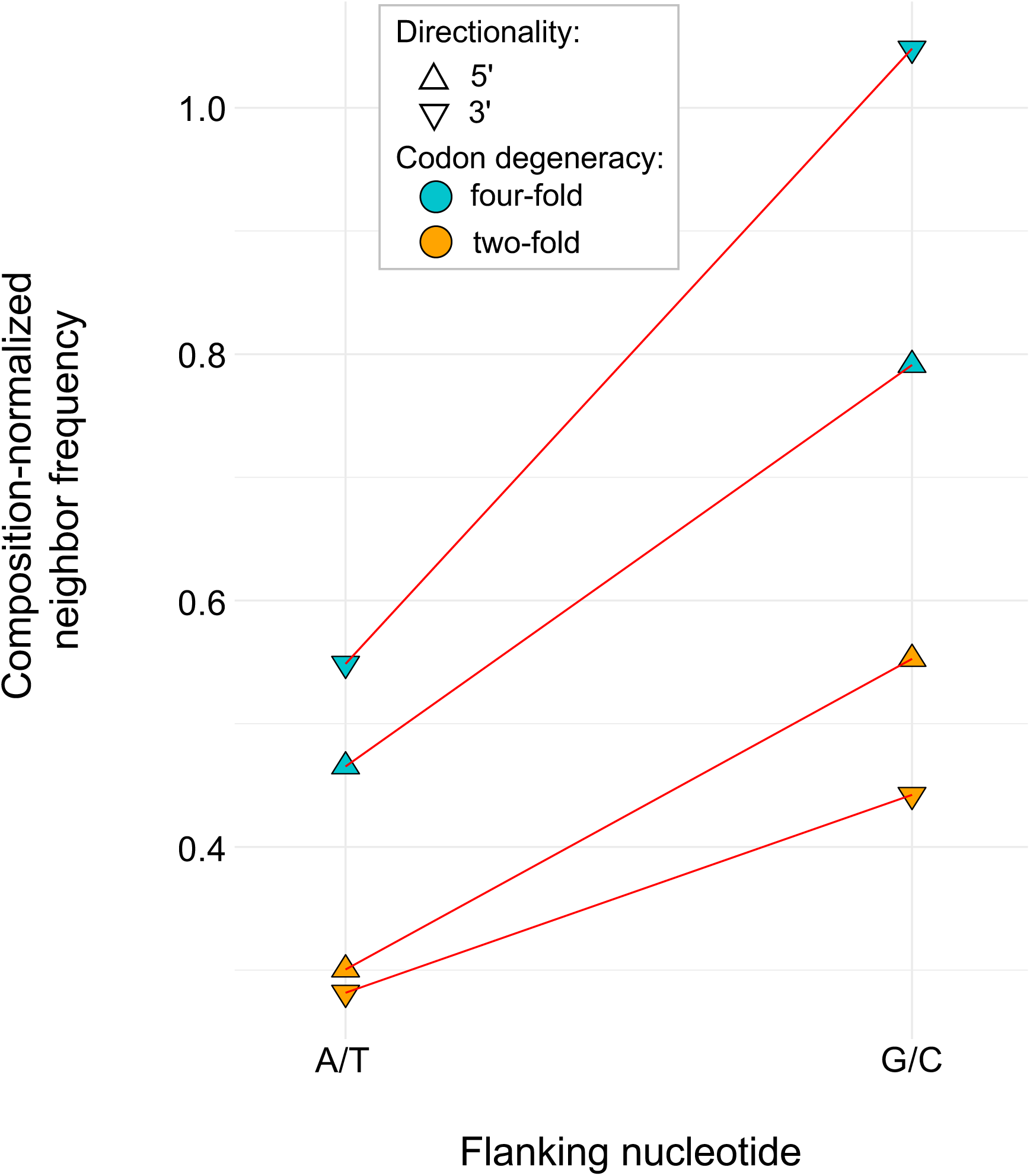
Frequency of nucleotides flanking synonymous polymorphisms at four-fold and two-fold degenerate sites normalized by base composition.

### The distribution of synonymous polymorphisms depends on their location within genes

There was no relationship between distance from the mtDNA control region and synonymous polymorphism. However, the combined data from all 12 protein-coding genes shows that the 5′ and 3′ ends of protein-coding genes contained fewer synonymous polymorphisms than centrally located sites within the open-reading frame. For example, the synonymous polymorphism count is two and five at the first and second 9-bp window (at the 5′ end), respectively (Fig. 7a). The last three 9-bp windows at the 3′ end averaged ∼6.7 synonymous polymorphisms per window (Fig. 7b). In contrast, centrally located 9-bp windows within the open-reading frame averaged ∼20.3 synonymous polymorphisms per windows (from the fourth to the sixth 9-bp window) (Fig. 7a). Similarly, A+T-content decreases along the same axis with increasing distance from both the start and stop codons (Figs. 7c and d).

**Fig. 7.**
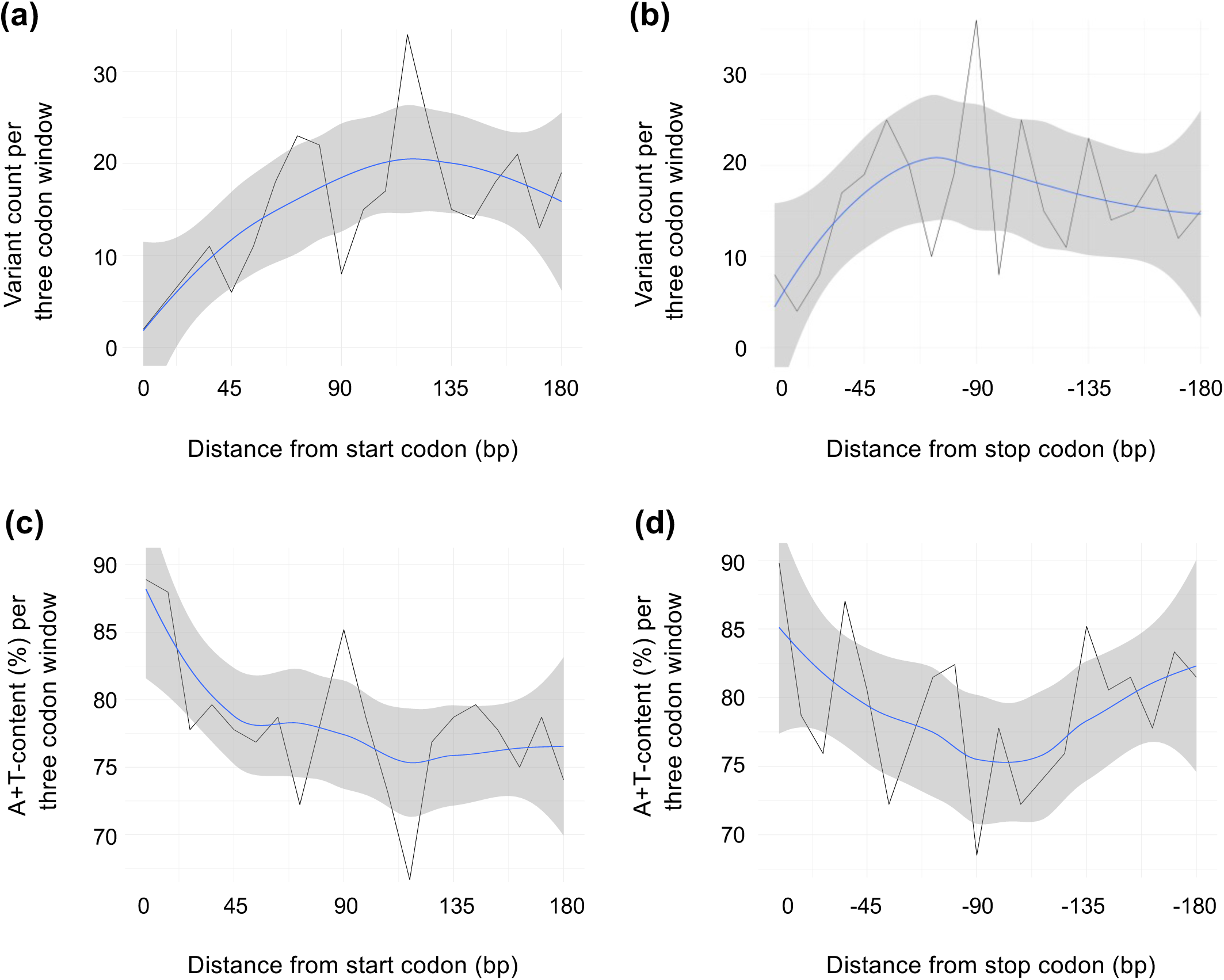
Number of synonymous polymorphisms and A+T-content in relation to the distance from a gene’s start and stop codons. a) Synonymous polymorphism as a function of distance from start codons. b) Synonymous polymorphism as a function of distance from stop codons. c) A+T-content as a function of distance from start codons. d) A+T-content as a function of distance from stop codons. The metrics were compiled across the 12 protein-coding genes for the first and last 180 bp with the blue line representing a LOESS curve fitted to the data with a 95% CI shown in grey.

### Small indel variants detected at low heteroplasmic frequencies with a clustered distribution in several hotspots in the mtDNA genome

There are two hypervariable mtDNA regions in the *C. elegans* natural isolates harboring small indels, namely (i) the noncoding region containing the D-loop, and (ii) a noncoding poly-A sequence between *atp6* and *tRNA_Lys_*. Exempting these two regions, 88 polymorphic small indels were identified. The vast majority of these indels were either a 1 bp insertion or a deletion and only two variants were 2 bp indels (an insertion and deletion each). Indel polymorphisms were found at one in every 22 tRNA and 350 protein-coding positions. It is worth noting that no indels were found in rRNA genes (supplementary file S2). In total, 29 (∼33%) small indels were identified in protein-coding genes, corresponding to seven deletions and 22 insertions. All but two of these 29 indels in protein-coding genes are heteroplasmic frameshift mutations, ranging in heteroplasmic frequency from 11% to 52%. Two indels were homoplasmic within the same isolate (XZ1516), a 1 bp deletion and a 1 bp insertion five nucleotides apart in the *nd2* gene, thereby preserving the reading frame. The tRNA genes harbored 56 (∼64%) small indel polymorphisms, corresponding to 47 single bp deletions, a single 2 bp deletion and eight 1 bp insertions. Apart from the two hypervariable regions, three (∼3%) indel polymorphisms were located in noncoding regions.

In addition to the hypervariable regions mentioned above, we identified additional small indel hotspots in four genes, *tRNA_Pro_, tRNA_Tyr_, nd4* and *nd5* (supplementary file S2). A *tRNA_Pro_* sequence corresponding to mtDNA:14-22 comprised of (A)_2_-T-(A)_5_ in the reference genome contained 32 and two cases of single A deletions and insertions, respectively. Natural isolate ECA740 incurred two A insertions within this region, with one insertion inferred in an ancestral reconstructed sequence and a subsequent second insertion leading to ECA740 having a double A insertion relative to the reference genome. The second hotspot occurs at position mtDNA:1,750 located within *tRNA_Tyr_* with 10 independent instances of an inferred T deletion and one T insertion. The secondary structure of these two tRNAs appears to be conserved. The third hotspot is found within a (T)_8_ homopolymeric run in *nd4* at position mtDNA:6,752. This site has three instances of a single T insertion and one case of a single T deletion. All four frameshift variants were found in low heteroplasmic frequencies ranging from 12−20%. The fourth and final indel hotspot occurs towards the 5′ end of *nd5* within a (T)_8_ homopolymeric sequence spanning positions mtDNA:11,722-11,729. We identified 11 cases of a single T insertion within this hotspot resulting in frameshifts spanning most of the *nd5* coding sequence. Similar to the *nd4* variants, these *nd5* frameshift variants were found in low heteroplasmic frequencies ranging from 11% to 25%. While insertions outnumbered deletions overall, the insertion/deletion bias is strikingly different between the tRNA and protein-coding genes. In tRNA genes, deletions were found to outnumber insertions by a factor of six. Conversely, insertions outnumbered deletions by a factor of three in protein-coding genes.

### First identification of novel heteroplasmic large mtDNA deletions and duplications in natural isolates of C. elegans

We identified eight heteroplasmic large structural variants (SVs), with each SV identified by three SV callers (Delly, Manta and Pindel). No variants (post-filtering) were called by only two of the callers employed. Of the eight variants, six were found to be large deletions while two were found to be duplications spanning the control region and neighboring genes (Table 2; Fig. 8; supplementary figs. S2 and S3). The heteroplasmic variant frequencies determined by read-depth analyses ranged from a low 4% up to 79% (Table 2). The two duplication variants found in strains PS2025 and MY23, while strikingly similar with respect to both location and duplication span, did not share any notable SNPs and do not appear to have the same origin. In particular, the precise breakpoint locations differed between the duplications.

**Fig. 8.**
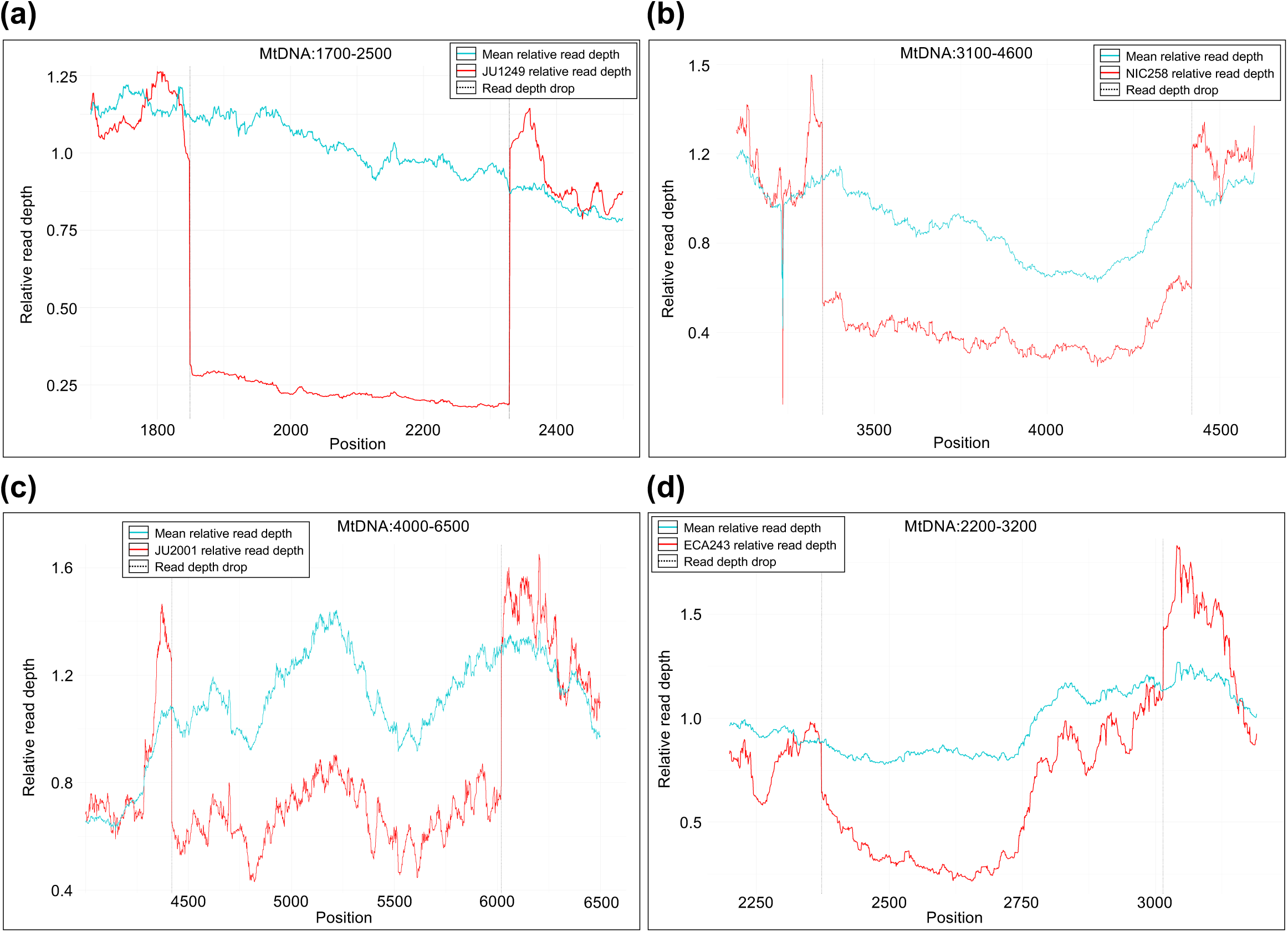
The distribution of read depth for the four mitochondrial genomes with the highest heteroplasmic frequencies of mtDNA deletions among the natural isolates. The horizontal axis represents the position in the reference strain mtDNA molecule and the vertical axis represents the relative read depth at any given position. The red and blue line are the distribution of relative read depth in the mtDNA harboring the deletion and mtDNA in all other natural isolates, respectively. The two grey vertical lines indicate the position of the deletion breakpoints. a) Deletion in natural isolate JU1249. b) Deletion in natural isolate NIC258. c) Deletion in natural isolate JU2001. d) Deletion in natural isolate ECA243.

**Table 2.** Large deletions identified by three SV callers (Delly, Manta and Pindel). The heteroplasmic frequencies were computed separately based on the ratio of the relative read depth of the deleted region in the focal strain relative to the read depth of the corresponding region across all other strains.

| Strain | Position | Type | Size (bp) | Genes affected | Heteroplasmic frequency |
| --- | --- | --- | --- | --- | --- |
| ECA243 | mtDNA:2,365-3,015 | Deletion | 651 | *nd1, atp6* | 0.41 |
| JU1249 | mtDNA:1,839-2,329 | Deletion | 491 | *nd1* | 0.79 |
| JU2001 | mtDNA:4,412-6,018 | Deletion | 1,607 | *tRNA<sub>Gln</sub>, tRNA<sub>Phe</sub>, ctb1, tRNA<sub>Leu</sub>, cox3* | 0.45 |
| NIC258 | mtDNA:3,341-4,419 | Deletion | 1,079 | *tRNA<sub>Leu</sub>, tRNA<sub>Ser</sub>, nd2, tRNA<sub>Ile</sub>, tRNA<sub>Arg</sub>, tRNA<sub>Gln</sub>* | 0.55 |
| QG2075 | mtDNA:9,797-11,815 | Deletion | 2,019 | *cox2, tRNA<sub>His</sub>, 16S rRNA, nd3, nd5* | 0.04 |
| QG2932 | mtDNA:8,327-12,320 | Deletion | 3,994 | *cox1, tRNA<sub>Cys</sub>, tRNA<sub>Met</sub>, tRNA<sub>Asp</sub>, tRNA<sub>Gly</sub>, cox2, tRNA<sub>His</sub>, 16S rRNA, nd3, nd5* | 0.095 |
| MY23 | mtDNA:12,024-452 | Duplication | 2,222 | *nd5, tRNA<sub>Ala</sub>, D-loop, tRNA<sub>Pro</sub>, tRNA<sub>Val</sub>, nd6* | 0.10 |
| PS2025 | mtDNA:12,015-444 | Duplication | 2,223 | *nd5, tRNA<sub>Ala</sub>, D-loop, tRNA<sub>Pro</sub>, tRNA<sub>Val</sub>, nd6* | 0.54 |

### Signature of purifying selection in heteroplasmic variants

Variants ranging in frequency from 3% to 97% within a natural isolate were characterized as heteroplasmic. In addition to the large heteroplasmic structural variants discussed above, of the 2,464 base substitutions and small indels variants in the natural isolates, only 6.6% (162) were heteroplasmic (Fig. 9a). Of these, 144 (∼89%) were within protein-coding genes. While only 4% and 11% of synonymous and nonsynonymous base substitutions were heteroplasmic, 100% of the small indel frameshift variants in the protein-coding genes were found to be heteroplasmic. The variant frequency distribution of small indel mutations differs between protein-coding and tRNA genes (Fig. 9b). The vast majority of small indels occurring in protein-coding genes are found at low heteroplasmic frequencies. In contrast, small indel mutations occurring in tRNA genes tend to have significantly higher frequencies (fixed or near fixation) (*U* = 90, *z* = 6.6878, *p* < 0.00001). The frequency distribution of heteroplasmic mtDNA mutations in the natural isolates is bimodal (Fig. 9a) and differs from the more uniform distribution observed for their counterparts in a long-term *C. elegans* MA experiment under minimal selection (Konrad et al. 2017; Fig. 9c). In protein-coding genes, the median frequency of heteroplasmic synonymous variants is significantly greater than that of heteroplasmic nonsynonymous variants (*U* = 1135, *z* = 2.9829, *p* = 0.00288). However, the median frequency of heteroplasmic nonsynonymous variants is not significantly greater than the median frequency of heteroplasmic frameshift variants (*U* = 534, *z* = 0.7614, *p* = 0.4473).

**Fig. 9.**
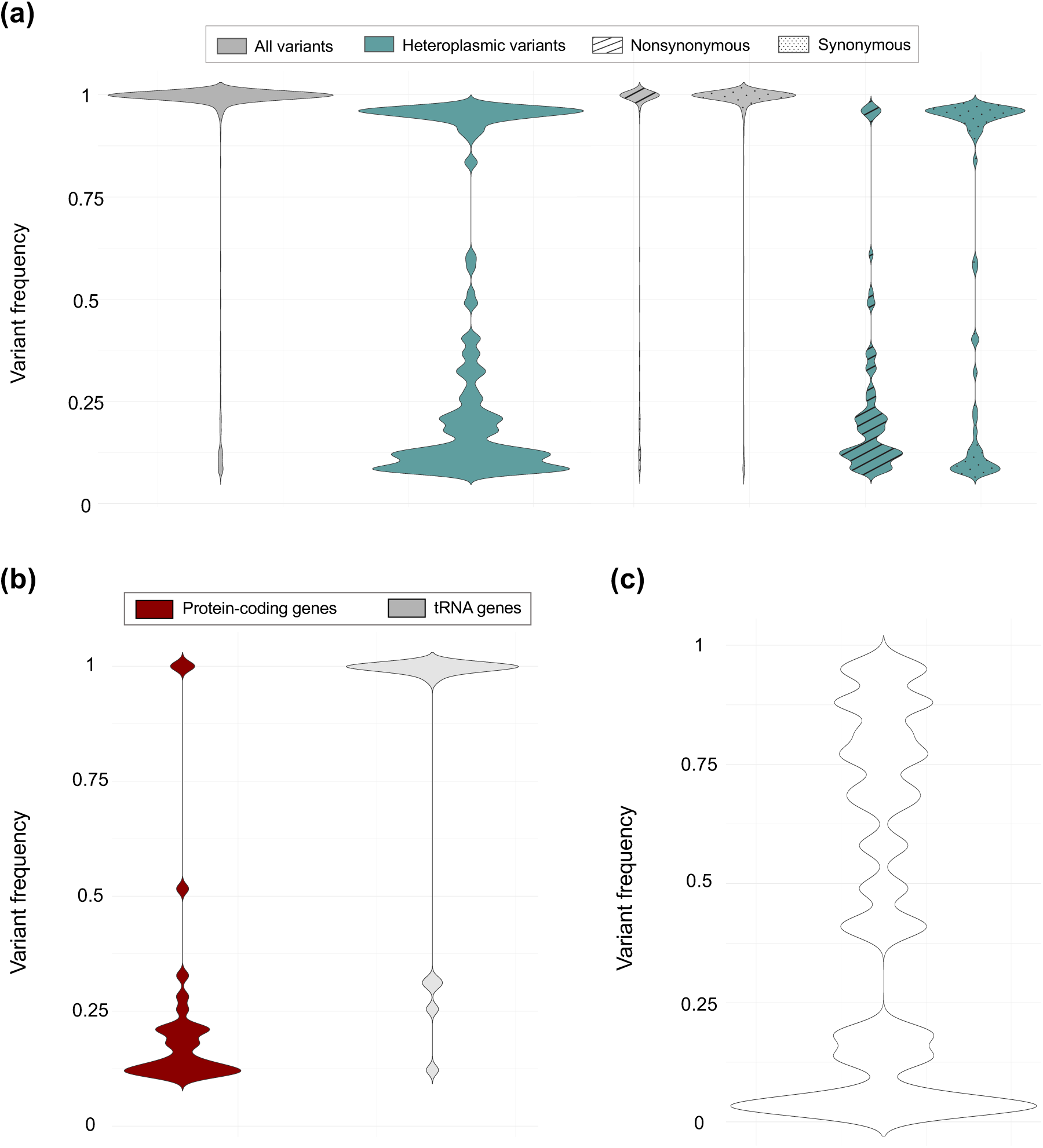
Variant frequency distribution for a) all mutations identified in the natural isolates comprising this study, b) all small indel variants identified in the natural isolates comprising this study, and c) all variants identified in a long-term *C. elegans* mutation accumulation experiment over 409 consecutive generations (adapted from Konrad et al. 2017). Variants with frequencies ranging from 0.03 to 0.97 were designated as heteroplasmic variants.

### McDonald-Kreitman tests detect no evidence of positive natural selection

To complement the metrics on mtDNA polymorphism and heteroplasmy in the natural isolates, we conducted McDonald-Kreitman tests (McDonald and Kreitman 1991) using *Caenorhabditis brenneri* as an outgroup species to investigate what proportion, if any, of the substitutions in the mtDNA protein-coding genes of the natural isolates are driven by positive adaptive evolution. We compared the within-species (*C. elegans* wild isolates) synonymous and nonsynonymous polymorphism to between-species synonymous and nonsynonymous divergence (supplementary table S2). The neutrality index across the 12 concatenated protein-coding genes was 0.79 (Fisher’s exact test: *p* = 0.11), which is not indicative of positive natural selection in the coding sequence between the two species. The neutrality index values fluctuate considerably between the different mtDNA genes analyzed, ranging from 0.42 to 3.3, but none of the genes show an excess of nonsynonymous changes between species (supplementary table S2).

### Codon usage bias has the potential to skew compositional and strand biases in the C. elegans mitochondrial genome

If the base composition of four-fold degenerate codon positions is strictly a function of mutation pressure, and not natural selection on synonymous mutations or specific local sequence context, these sites should have similar base composition between codon families. Four of the seven four-fold degenerate codon families deviated significantly in their nucleotide distribution at their third base position from other four-fold degenerate codon families (supplementary table S3). Furthermore, five of the 13 two-fold degenerate codon families also show significant variation in their nucleotide distribution at their 3^rd^ base position (supplementary tables S4 and S5). The most commonly used codons in each codon family (preferred codons) contain either a T (most common) or an A in their 3^rd^ base position. The codon usage preferences in *C. elegans* mtDNA suggest natural selection for codon usage, which in addition to the mutational spectrum, introduces a further base compositional bias and strand asymmetry to the *C. elegans* mitochondrial genome. A comparison of synonymous mutations present only in single mitotypes (rare) versus variants that are shared by more than a single mitotype, lends further support for natural selection on codon preference in *C. elegans* mtDNA. In rare synonymous variants, mutations from preferred codons outnumber mutations to preferred codons by 357 to 245. In contrast, in variants that are shared by more than a single mitotype, mutations from and to preferred codons are similar (364 and 333, respectively). This difference between rare and shared synonymous variants is significant (*G* test-statistic = 6.56, *d.f.* = 1, *p* = 0.0104). Rare variants are thought to be more representative of the mutation spectrum and less affected by natural selection than more common (shared) variants. The greater relative abundance of mutations to preferred synonymous codons in shared variants compared to rare variants is consistent with natural selection for preferred synonymous codons.

## Discussion

### Natural populations have fewer mtDNA mutations typically associated with oxidative damage relative to laboratory populations

Polarized DNA polymorphism from natural isolates was used to test if the mtDNA mutational patterns observed in laboratory experiments recapture the mutational spectrum of natural populations. Laboratory experiments exhibit varying patterns but still converge on high proportions of G/C → T/A transversions, which have been interpreted as signatures of oxidative damage (Waneka et al. 2021; Leuthner et al. 2022). This hypothesis has been reinforced by observations of mutational strand bias (Cheng et al. 1992; Waneka et al. 2021). One of the challenges when comparing natural populations to laboratory experiments (even more so when these laboratory lines are the result of several generations of mutation accumulation under minimal selection) is the difference in the strength of purifying selection, which could have significant effects on the observed mutational spectrum. To mitigate the effect of selection on the observed mutational spectrum in the natural isolates, we focused on sites where selection is expected to be at a minimum, such as four-fold degenerate sites. Furthermore, in the natural isolates’ dataset, we compared the results between (i) polymorphisms where polarity was inferred using ancestral reconstruction, and (ii) rare variants that were only observed in single mitotypes. Purifying selection is a potent force acting on mtDNA polymorphism in the natural isolates. This is evident in the distribution of mtDNA substitutions which are strongly associated with codon position and site degeneracy. Four-fold degenerate sites harbor the vast majority of mutations in the natural isolates. Furthermore, the high *Ts*/*Tv* ratio at two-fold degenerate sites compared to four-fold degenerate sites is a prime example of how strongly nonsynonymous mutations are counter-selected in the wild.

Within transitions and transversions, there are similarities in the ranking of proportions of different base substitutions between the natural isolates and laboratory experiments (Waneka et al. 2021; Leuthner et al. 2022). G/C → A/T transitions are more frequent than the reverse transition of A/T → G/C, contributing to an A+T-mutational bias in both the natural isolates and laboratory experiments. The main source of transversions in both natural isolates and laboratory lines are G/C → T/A mutations, further adding to the mtDNA A+T-mutational bias. However, the natural isolates exhibit a significantly greater *Ts*/*Tv* ratio than the laboratory populations. The main contributor to the difference in the *Ts*/*Tv* ratio between the mutation spectra of the natural isolates *vs.* laboratory populations is a far greater proportion of G/C → T/A transversions in the laboratory experiments (Waneka et al. 2021; Leuthner et al. 2022). The proportion of G/C → T/A transversions is approximately 4× greater at four-fold degenerate sites in the laboratory experiments relative to the natural isolates. G/C → T/A transversions have been associated with oxidative damage of DNA (Cheng et al. 1992; Kirkwood and Kowald 2012), and the striking difference in their proportions between laboratory experiments and natural populations might suggest that laboratory grown populations are more prone to mtDNA oxidative damage.

Metabolic rate has been suggested to be a major determinant of mtDNA evolution (Martin and Palumbi 1993; Rand 1994; Gillooly et al. 2005). The rate-of-living theory (Rubner 1908; Pearl 1928) and the free radical theory (Harman 1956) that expands upon it, proposed that higher metabolic rate organisms have shorter life spans due to the faster accumulation of damages from metabolic by-products and more specifically due to the mutagenic nature of ROS species (Beckman and Ames 1998). Moreover, mtDNA maybe especially prone to oxidative damage compared to nuclear DNA (Richter et al. 1988). However, the relationships between basal metabolic rate and the rates of mutations and molecular evolution in mtDNA have remained somewhat murky and adaptation associated with other life-history traits such as life span may have been more important (Galtier et al. 2009). Many attributes of the base composition and strand bias in mitochondrial genomes have been ascribed to DNA damage by ROS.

*C. elegans* is a widely used laboratory model and has been genetically modified through decades of domestication to adapt to cultivation methods enforced upon it (Frézal and Félix 2015). In the wild, *C. elegans* lives through boom-and-bust cycles depending on temperature and food supplies, entering a diapause state called dauer through periods of insufficient resources. The dauer larval state has been found to increase resistance to oxidative damage, leading to increased life-span (Larsen 1993). The concentration of oxygen on petri dishes is much higher than that in the preferred habitat of wild *C. elegans* and during its domestication, laboratory strains have lost many hyperoxia avoidance behaviors that are present in natural isolates (Chang et al. 2006). We hypothesize that laboratory growth conditions for *C. elegans*, with unlimited food, relatively unrestricted space in the case of MA experiments, and an absence of the multitude of environmental stressors present in the wild lead to higher metabolic rates and greater oxidative damage over the course of a laboratory experiment than that usually experienced in the wild.

### Number of polymorphic base substitutions varies by gene and ETC complex

Purifying selection is the main evolutionary force acting on mtDNA polymorphism in *C. elegans* natural isolates. The distribution of polymorphism is strongly linked to site degeneracy, as shown, for example, by the high *Ts/Tv* at two-fold compared to four-fold degenerate sites. However, the strength of purifying selection may not be homogeneously distributed across all mtDNA protein-coding genes. ETC complex I genes have higher nonsynonymous to synonymous polymorphism ratio (*N*:*S*), most apparent in *nd6* and *nd2.* Indeed, five of the seven mtDNA genes in this complex exhibit higher than average *N*:*S* ratios. Conversely, genes in complex IV tend to have a lower N:S, suggestive of strong purifying selection. These patterns of mtDNA polymorphism are concordant with an analysis of mtDNA divergence between *Caenorhabditis* species which found the highest and lowest *dN/dS* in ETC complex I and IV genes, respectively (Nabholz et al. 2013). Furthermore, this study found an inversely proportional relationship between *dN/dS* and mtDNA transcript abundance of mitochondrial-encoded OXPHOS genes. Although the results were based on transcript rather than protein abundance, this negative relationship is consistent with the hypothesis that highly expressed proteins have slower rates of evolution (Drummond et al. 2005). However, an increasing number of studies have associated mtDNA mutations in ETC complex I with life-history evolution (Garvin et al. 2015; Camus et al. 2017; Li et al. 2024), or have identified ETC complex I as a target of directional selection (Mishmar et al. 2006). For example, different mitotypes distinguished primarily by ETC complex I mutations are associated with thermal adaptation in *Drosophila melanogaster* (Camus et al. 2017). Similarly, in the agricultural pest *Spodoptera frugiperda*, the “rice” mitotype at the forefront of pest invasions differed from the “corn” mitotype by 18 nonsynonymous mutations, 14 of which are in ETC complex I (Li et al. 2024). ETC complex I is often associated with positive selection and appears to be one of the primary targets of local adaptation in mitochondrial genomes (Mishmar et al. 2006; Garvin et al. 2015; Camus et al. 2017; Li et al. 2024).

### The effect of A+T-content of flanking nucleotides on synonymous polymorphism

There is a significant difference between A/T versus G/C neighbors on synonymous polymorphism. The probability of a site exhibiting synonymous polymorphism decreases with A+T-content of its adjacent nucleotides. Whereas variation in nonsynonymous polymorphism between genes is frequently explained by differences in the relative strength of natural selection, synonymous polymorphism is often presumed to be neutral. Population-genetic processes such as genetic hitchhiking or background selection (Maynard Smith and Haigh 1974; Charlesworth et al. 1993) which can introduce variation in synonymous polymorphism in genomes undergoing genetic recombination, should not contribute to between-gene variation in synonymous polymorphism in completely linked genomes such as the *C. elegans* mitochondrial genome. There is a potential for selection on synonymous mutations through codon usage and mRNA secondary structure, which can vary among genes. However, local sequence context appears to have a significant contribution to between-gene variation in synonymous polymorphism as demonstrated by a correlation between A+T-content in the 1^st^ and 2^nd^ codon positions and synonymous polymorphism, usually in 3^rd^ codon position.

The effect of local sequence context also extends to within-gene variation in synonymous polymorphism. The 5′ ends of *C. elegans* mtDNA genes, and 3′ ends to some extent as well, exhibit less synonymous polymorphism than centrally located sites. This coincides with a higher A+T-content at the 5′ and 3′ ends across the 12 mtDNA protein-coding genes. Does this within-gene variation in base composition and mutation rate/synonymous polymorphism indicate some structural constraints on mRNA structure? A possible neutral explanation for this pattern is that amino acid composition is influenced by mutation pressure in such a manner that the consequences of high A+T mutation pressure are not just confined to synonymous sites but nonsynonymous sites as well (Foster et al. 1997). Nonsynonymous sites within genes that are less important for optimal protein function might therefore be more affected by A+T mutation pressure and code for amino acids with high A+T-content at their nonsynonymous sites, usually at the 1^st^ and 2^nd^ codon positions. Consequently, because of the influence of local sequence context on the frequency of mutations, the probability of synonymous polymorphism would be lower in these high A+T-regions of genes. However, it is also possible that this pattern of high A+T-content at the 5′ end of genes shares functional similarities to many prokaryotic genomes that also have higher A+T-content at their genic 5′ ends (Allert et al. 2010; Tuller et al. 2010). In *E. coli*, protein expression levels have been shown to be dependent on high A+U-content and low secondary structure of mRNA, particularly at the gene’s 5′ region, and to a lesser extent, on high A+U-content in the 3′ region (Allert et al. 2010; Goodman et al. 2013). Considering the prokaryotic origin of mitochondria, perhaps selection to maintain (i) high A+T-content, and/or (ii) low complexity to prevent mRNA secondary structures that might interfere with optimal translation efficiency is still important in mitochondrial genomes.

### Natural selection on codon usage bias

Unequal use of synonymous codons can stem from compositional and strand biases caused by the mutational spectrum and natural selection for particular codons within a family of synonymous codons (Hershberg and Petrov 2008). The observed deviation in base composition and strand symmetry at four-fold degenerate sites from the expected equilibrium value based on the mutation spectrum in these *C. elegans* natural isolates could be explained by codon usage selection. Furthermore, natural selection for optimal codon usage could potentially influence the frequency spectrum of polymorphisms at degenerate sites. Families of synonymous codons in the *C. elegans* mitochondrial genome always favor an A or T at degenerate sites in the coding sequence, which is expected given the prevailing G/C → A/T mutational bias. Furthermore, most codon families with four-fold degenerate sites are also biased towards having a T instead of an A in their 3^rd^ base position, which is also expected given the observed compositional and mutational strand asymmetry. An example of a strong preference for a T is the family of arginine codons as 29 of 31 arginine codons in the *C. elegance* reference mtDNA contain a T in their 3^rd^ base position. However, some four-fold degenerate codon families are biased towards an A in their 3^rd^ base position which runs counter to the compositional and mutational strand asymmetry suggesting a role for natural selection for codons ending in A in these families. An example of a strong preference for A at 3^rd^ base position is the family of serine codons that start with AG. In this family, codon AGA is present 126 times compared to 61 instances of codon AGT in the reference genome. If mutational bias alone is responsible for codon usage bias, we expect that four-fold degenerate codon families have the same distribution of bases in their 3^rd^ base position, which they do not. Furthermore, the significantly higher proportion of mutations to preferred codons among inferred ancestral variants compared to rare variants is consistent with natural selection for preferred codons. The spectrum of synonymous polymorphism in *C. elegans* mtDNA is therefore a function of mutational patterns, which include base compositional biases, strand asymmetry and local sequence context, as well as natural selection for specific synonymous codons.

### First identification of novel heteroplasmic structural variants in wild populations of C. elegans

Large structural mtDNA variants have previously only been described for laboratory populations of *C. elegans* (Tsang and Lemire 2002; Liau et al. 2007; Dubie et al. 2020, 2024; Sequeira et al. 2024). The natural isolates analyzed here add eight new heteroplasmic structural variants comprising six deletions and two duplications. As all of the mtDNA genes are essential, genic mtDNA deletions are always heteroplasmic and appear to be maintained in populations by the balance between replication or transmission advantage and fitness cost (Liau et al. 2007; Gitschlag 2016, 2020; Dubie et al. 2020, 2024; Sequeira et al. 2024). The six deletions described here affect nine protein-coding genes and multiple tRNA genes. Notably, regions in the *C. elegans* mtDNA have not been found to be deleted in either laboratory or natural populations include the first few genes downstream of the D-loop (*nd6*, *nd4L*, *tRNA_Pro_*_,*Val*,*Trp*,*Glu*_, and 12S rRNA) and a noncoding region between *nd4* and *cox1* which has been suggested to harbor a second replication origin. These regions that are not known to have been affected by deletions are possibly essential for mtDNA maintenance. In addition to the six deletions, duplications encompassing the D-loop were identified in two natural isolates. Duplications of the mtDNA replication origin can possibly be favored in intra-individual selection by providing additional replication initiation targets (Blanc and Dujon 1980). However, some studies have suggested that mtDNA structural variants, including duplications, can be a conduit for molecular adaptation (Minhas et al. 2023).

### Conclusions

Polarized polymorphisms in *C. elegans* natural isolates allowed us to study a larger sample of mtDNA mutations than previously achieved in laboratory settings (Denver et al 2001; Konrad et al 2017; Waneka et al 2021; Leuthner et al 2022). While this approach may lack the controls possible in laboratory experiments, we show that the laboratory environment can introduce a unique set of biases to the mutational spectrum of mtDNA, possibly due to higher levels of mtDNA oxidative damage in experimental populations. ETC complex I genes have greater levels of nonsynonymous polymorphism than other protein-coding mtDNA genes in *C. elegans*. Whereas this could suggest less stringent selection on ETC complex I genes, several studies have linked complex I mutations with local adaptation (Mishmar et al. 2006; Garvin et al. 2015; Camus et al. 2017; Li et al. 2024). Local sequence context, especially A+T-content of flanking nucleotides, significantly influences synonymous polymorphism between- and within-genes. Comparisons between rare polymorphisms found only in single mitotypes and inferred ancestral polymorphism present in more than a single mitotype suggest that natural selection contributes to codon usage bias in *C. elegans* mitochondrial genomes. Finally, we identify the first large heteroplasmic mtDNA structural variants in natural isolates of *C. elegans*. Large, and likely highly disruptive, these structural variants in wild populations possibly possess selfish properties akin to those observed in similar variants arising in experimental laboratory populations. Inferences of mutation spectra using data from natural populations is not without caveats, particularly regarding the possible effects of selection on synonymous sites and possible errors in inferring ancestral states. Nevertheless, the study of mutation spectra in natural isolates is highly relevant for the interpretation of molecular evolution data, especially in organisms such as *C. elegans* whose ecological habitat and environment differ significantly their rearing conditions in laboratories.

## Materials and Methods

### C. elegans natural isolates used in this study

Genomic DNA sequences of 550 isotypes corresponding to 1,524 natural isolates of *C. elegans* were downloaded from the CaeNDR database (Crombie et al. 2024). A complete list of strains used in this study is provided in supplementary file S1. Reads mapping to the mitochondrial genomes were retained from the downloaded alignment files of the 550 isotypes. The reference mitochondrial genomes of two congeneric species, *C. brenneri* and *C. tropicalis,* (Yang et al. 2016) were downloaded from NCBI for use as outgroups. During a preliminary analysis, we identified 19 lines (supplementary fig. S1) exhibiting an unusual pattern of heteroplasmic variant frequencies. This pattern comprised of several parallel variants (shared with other isotypes) in very similar heteroplasmic frequencies. The combination of multiple parallel mutations in similar frequencies within a single isotype led us to consider contamination from another isotype as the most likely explanation. The mtDNA sequences of these 19 lines/isotypes were removed from further analysis, resulting in a final set of 531 isotypes.

### Phylogenetic reconstruction

A consensus sequence for each natural isolate’s alignment data was built using Samtools (Li et al. 2009). The mtDNA consensus sequences for all 531 isotypes and the outgroup reference sequences were aligned using MAFFT (Katoh and Standley 2013). All sequences were trimmed at two locations using SEQKIT (Shen et al. 2016). The first trim removed a variable homopolymeric run between *atp6* and *tRNA_Lys_*. The second trim removed a highly variable noncoding sequence between *tRNA_Ala_* and *tRNA_Pro_* thought to contain the origin of replication. Identical mitochondrial consensus sequences (100% match) were clustered using CD-HIT (Li et al. 2001) and one sequence for each cluster was retained for further analysis, reducing the original number of 531 isotype sequences to 347 unique mitotypes. A maximum likelihood tree was built with the 347 unique mitotype sequences and the two reference sequences from the other *Caenorhabditis* species using IQ-TREE2 (Minh et al. 2020) with TIM2 + F + I + R4 as suggested by Model Finder Plus (Kalyaanamoorthy et al. 2017). The ML phylogenetic tree was used to identify the most basal strains or subtree of natural isolates which comprised six strains from Kauai Island in the Hawaiian archipelago. The strain XZ1516 from this basal group was used as an outgroup for subsequent analysis of the natural isolates. A second maximum likelihood tree was subsequently built with TN + F + I + R4 as suggested by Model Finder Plus (Kalyaanamoorthy et al. 2017) using XZ1516 as an outgroup.

### Ancestral Sequence Reconstruction (ASR) and inference of the ancestral versus derived variant state

We used ARPIP (Jowkar et al. 2022) to reconstruct the ancestral sequence for each node in the phylogenetic tree. A custom R script using SEQINR (Charif and Lobry 2007) was used to catalog the ancestral mutations inferred by ARPIP along the phylogenetic tree by comparing each node to its two descendants and iterating this pattern until a leaf was reached. This method can take into account parallel mutations and only counts mutations shared by a clade as a single event. To polarize the variants found in the natural isolates, the variants of each strain were compared to their most immediate ancestors for which a reference had been previously reconstructed with ARPIP. To call the variants, the BAM files downloaded from the CaeNDR were first converted back to FASTQ with SAMtools (Li et al. 2009), the reads were aligned to the reconstructed ancestor’s sequence using BWA (Li and Durbin 2009), and optical duplicates were marked using Picard MarkDuplicates (Van der Auwera and O’Connor 2020). Finally, the variant calling itself was performed using GATK HaplotypeCaller (Van der Auwera and O’Connor 2020) and variant frequency information was added using BCFTOOLS (Li 2011). Subsequent variant annotation was performed in a custom script. For variant annotation, the start and stop positions for each reference’s genes were determined using BLAST (Camacho et al. 2009) against the N2 reference sequence. A custom R script using the VCFR (Knaus and Grünwald 2017) and SEQINR (Charif and Lobry 2007) packages was used to annotate the variants. There were several instances of a cluster of lines descended from the same ancestor that had a number of shared variants originally identified as independent parallel mutations. In these instances, we only retained a single representative mitotype with the highest number of mutations from the cluster.

### Association between flanking nucleotides and mutations

The frequencies of the 5′ and 3′ immediate neighbors of synonymous polymorphisms were analyzed to assess a potential mutational bias associated with flanking nucleotides. The neighboring bases of synonymous mutations were analyzed separately for four-fold degenerate and two-fold degenerate sites because codons with four-fold and two-fold degenerate sites in the 3^rd^ base position are more likely to have a G/C and A/T in their 2^nd^ base position, respectively. The counts of bases flanking synonymous mutations at two-fold or four-fold degenerate sites were normalized by the base composition of 5′ or 3′ nucleotides flanking all two-fold or four-fold degenerate sites in the reference mtDNA. The observed counts were then statistically compared to the expected counts based on the base composition.

### Distribution of synonymous mutations and A+T-content within mtDNA genes

In order to analyze the distribution of synonymous mutations within the 12 protein-coding mtDNA genes, a sliding window analysis (nonoverlapping three codon windows) was performed on both cumulative synonymous mutation counts and A+T-content within genes.

### Codon Usage Bias

Four-fold degenerate sites were used to draw conclusions about the mutation spectrum in natural populations of *C. elegans*. However, selection on codon usage in many organisms has the potential to bias the observed base substitution pattern. In order to investigate the possible effect of natural selection on the preferred use of certain codons that could lead to the observed bias in the base composition and mutation spectrum in the mitochondrial genome, we compared the observed proportion of each amino acid codon to its expected value based on the inferred mutation spectrum. We used the observed base composition at the third base position of codons to compute a RSCU accounting for compositional bias (Sharp et al. 1986). We first tested if the observed partition of codons for each amino acid with four-fold degenerate sites showed a significant deviation from the overall distribution of nucleotides at these sites. Post-hoc tests were performed comparing each amino acid’s third base composition to that of all other amino acids. For two-fold degenerate sites, the same analyses were performed but the A/G and T/C ending amino acids were analyzed separately. All tests of statistical significance were conducted using Pearson’s χ*^2^* test with 10,000 simulations and Bonferroni corrections.

The method used to polarize mutations resulted in two categories of polymorphisms, namely, (i) ancestral variants that were inferred between nodes in the reconstructed phylogeny, and (ii) rare variants which were only found in a single mitotype or strain, and inferred to have occurred between a node and a leaf in the phylogeny. Variants shared by large phylogenetic units, inferred to be ancestral, were expected to be, on average, older than variants only found in a single strain or mitotype. Conversely, rare variants that are only found in a single mitotype are expected to be, on average, younger, or perhaps less favored by natural selection than common variants (Zhu et al. 2017). We compared mutations to and from preferred synonymous codons between rare and ancestral synonymous polymorphisms to test for natural selection on synonymous codon usage. All amino acids were assigned their preferred codon as being the most used codon for that amino acid within the *C. elegans* N2 mtDNA reference genome (WS283). A *G*-test was performed to test for significance between ancestral and rare polymorphisms.

### Structural variants (SV)

Three SV callers were used to analyze the mtDNA genomes of the 531 isotypes, namely Manta (Chen et al. 2016), DELLY (Rausch et al. 2012) and Pindel (Ye et al. 2009) using the WS283 mitochondrial genome of *C. elegans* for reference. The variants were filtered to (i) remove variants falling within the region between *tRNA_Ala_* and *tRNA_Pro_*, a highly variable region, and (ii) remove variants supported by a low number of reads. This second filter was set at “PAIR_COUNT > 12” for the variants obtained through Manta, “PE >= 10 && SR >= 5” for the variants obtained with DELLY and “DP > 500 && VAF > 0.03” for the variants obtained with Pindel. The filtering itself was done using BCFtools (Li et al. 2011). Variants detected by two or more callers were visually confirmed in Integrated Genome Viewer (IGV) (Robinson et al. 2011) before further analysis. During preliminary analyses, two mtDNA structural variants were called as deletions spanning almost the entirety of the mitochondrial genome. Since these callers treat the genome as linear with artificial ends downstream of the D-loop rather than circular, duplications spanning the ends of the reference genome can lead to variants being incorrectly designated as deletions. In order to circumvent this limitation, a modified reference genome with its ends placed in a noncoding region between *nd4* and *cox1* at mtDNA:7,773 was used. For each variant, the heteroplasmic frequency was calculated as the ratio of the relative mean read depth of the variant (the mean read depth within the variant’s region relative to the mean read depth outside of the variant’s region) over the relative mean read depth of the region across all other lines without SVs. Per position read-depth was obtained using Samtools (Li et al. 2009). Due to less effective read-mapping in the highly variable D-loop region, reads between mtDNA:12,637-13,794 were not used in the variant frequency calculations. Variants spanning this region had their frequencies computed separately upstream and downstream within the duplicated region and averaged.

## Supporting information

Supplementary File S1. List of 1,524 C. elegans natural isolates used in this study whose genomic DNA sequences downloaded from CaeNDR.

Supplementary File S2. List and details of 2,464 mtDNA variants (88 small indels and 2,376 SNPs) analyzed in this study.

## Supplementary Material

Supplementary material is available at *Genome Biology and Evolution* online.

## Acknowledgements

The computations were enabled by resources provided by the National Academic Infrastructure for Supercomputing in Sweden (NAISS) and the Swedish National Infrastructure for Computing (SNIC) at Uppmax, [partially funded by the Swedish Research Council through grant agreements no. 2022-06725 and no. 2018-05973].

## Author Contributions

V.K. and U.B. conceived, designed and jointly supervised the study. A.S. conducted the bioinformatic phylogenetic analyses and generated the variant calls. A.S. and U.B. analyzed the data. All authors contributed to manuscript preparation.

## Data Availability

All data associated with this study have been made available in the supplementary materials.

