## Supplementary material for "Mutational Biases and Selection in Mitochondrial Genomes: Insights from a Comparative Analysis of Natural and Experimental Populations of *Caenorhabditis elegans*": Suppl Tables S1-S4, Suppl Figures S1-S3.

**Supplementary Table S1.** Comparisons of observed and expected counts of synonymous polymorphisms with flanking G/C or A/T bases. Synonymous polymorphisms were analyzed separately for two-fold and four-fold degenerate sites.

| Codon degeneracy | Neighbor direction | Neighboring nucleotide | Observed counts | Expected counts | *G*-test statistic | *p*-value |
|---|---|---|---|---|---|---|
| **Two-fold** | **5′** | **A/T** | 729 | 773.43 | 29.27 | $6.3 \times 10^{-8}$ ** |
| | | **G/C** | 105 | 60.57 | | |
| | **3′** | **A/T** | 566 | 640.80 | 34.91 | $3.5 \times 10^{-9}$ ** |
| | | **G/C** | 268 | 193.20 | | |
| **Four-fold** | **5′** | **A/T** | 161 | 243.28 | 41.54 | $1.2 \times 10^{-10}$ ** |
| | | **G/C** | 739 | 656.72 | | |
| | **3′** | **A/T** | 485 | 621.56 | 90.60 | $< 2.2 \times 10^{-16}$ ** |
| | | **G/C** | 415 | 278.44 | | |

**Supplementary Table S2.** McDonald-Kreitman test-statistics and associated significance level for each of the 12 protein-coding genes of the *C. elegans* mtDNA genome. The notations $d_N$ and $d_S$ represent fixed nonsynonymous and synonymous difference between *C. elegans* and *C. brenneri*, respectively, and $p_N$ and $p_S$ represent polymorphic nonsynonymous and synonymous changes within the *C. elegans* natural isolates.

| **Gene** | $d_S$ | $d_N$ | $p_S$ | $p_N$ | **Neutrality index** | ***p*-value; Fisher's exact test** |
|---|---|---|---|---|---|---|
| *nd6* | 44 | 23 | 31 | 11 | 0.68 | 0.40 |
| *nd4L* | 12 | 5 | 10 | 2 | 0.48 | 0.66 |
| *nd1* | 78 | 18 | 61 | 9 | 0.64 | 0.40 |
| *atp6* | 52 | 11 | 34 | 3 | 0.42 | 0.24 |
| *nd2* | 76 | 23 | 40 | 14 | 1.16 | 0.70 |
| *ctb1* | 93 | 17 | 72 | 10 | 0.76 | 0.68 |
| *cox3* | 75 | 1 | 46 | 2 | 3.26 | 0.56 |
| *nd4* | 113 | 14 | 72 | 12 | 1.35 | 0.52 |
| *cox1* | 121 | 7 | 117 | 5 | 0.74 | 0.77 |
| *cox2* | 55 | 5 | 44 | 4 | 1.00 | 1.00 |
| *nd3* | 27 | 4 | 17 | 2 | 0.79 | 1.00 |
| *nd5* | 143 | 21 | 81 | 9 | 0.76 | 0.55 |
| **Total** | 889 | 149 | 625 | 83 | 0.79 | 0.11 |

**Supplementary Table S3.** Comparison of the base distribution at four-fold degenerate sites between codon families. *Ad-hoc* tests were performed comparing the base composition at synonymous sites for individual codon families to four-fold degenerate sites in other codon families. The *p-value* threshold for *ad-hoc* tests is 0.05/7 = 0.0071. The Pearson's $\chi^2$ was calculated from 10,000 simulations because some bases at the 3rd codon position were present in fewer than five codons per amino acid.

| Amino Acid | T | A | G | C | Pearson's $\chi^2$ test statistic (*p*-value) |
|---|---|---|---|---|---|
| **Alanine** | 62 | 39 | 4 | 9 | 16.59 (0.0014*) |
| **Glycine** | 93 | 60 | 29 | 5 | 16.89 (0.0018*) |
| **Proline** | 27 | 39 | 8 | 7 | 8.24 (0.0399) |
| **Serine 1** | 61 | 126 | 39 | 6 | 57.98 (9.99e-05**) |
| **Serine 2** | 84 | 51 | 5 | 5 | 18.97 (0.0008*) |
| **Threonine** | 62 | 69 | 6 | 7 | 8.36 (0.0370) |
| **Valine** | 118 | 98 | 28 | 9 | 2.19 (0.5365) |
| **TOTAL** | 507 | 482 | 119 | 48 | 90.74 (9.99e-05**) |

**Supplementary Table S4.** Comparison of base distribution at two-fold degenerate sites between codon families ending in T or C bases. *Ad-hoc* tests were performed comparing the base composition at synonymous sites for individual codon families to two-fold degenerate sites in other codon families ending in T or C bases. The *p-value* threshold for *ad-hoc* tests is 0.05/7 = 0.0071. The Pearson's $\chi^2$ was calculated from 10,000 simulations because some bases at the 3rd codon position were present in fewer than five codons per amino acid.

| Amino Acid | T | C | Pearson's $\chi^2$ test statistic (*p*-value) |
|---|---|---|---|
| **Cysteine** | 45 | 2 | 1.37 (0.3170) |
| **Aspartic Acid** | 55 | 8 | 1.20 (0.3763) |
| **Phenylalanine** | 429 | 23 | 17.27 (9.99e-05*) |
| **Histidine** | 45 | 13 | 15.11 (0.0012*) |
| **Isoleucine** | 257 | 23 | 0.34 (0.6055) |
| **Asparagine** | 139 | 12 | 0.26 (0.6721) |
| **Tyrosine** | 144 | 29 | 19.95 (2e-04*) |
| **TOTAL** | 1,114 | 110 | 36.72 (9.99e-05**) |

**Supplementary Table S5.** Comparison of base distribution at two-fold degenerate sites between codon families ending in A or G bases. *Ad-hoc* tests were performed comparing the base composition at synonymous sites for individual codon families to two-fold degenerate sites in other codon families ending in A or G bases. The *p-value* threshold for *ad-hoc* tests is 0.05/6 = 0.0083. The Pearson's $\chi^2$ was calculated from 10,000 simulations because some bases at the 3rd codon position were present in fewer than five codons per amino acid.

| Amino Acid | A | G | Pearson's $\chi^2$ test statistic (*p*-value) |
|---|---|---|---|
| **STOP** | 10 | 2 | 1.35 (0.2362) |
| **Glutamic Acid** | 52 | 26 | 63.99 (9.99e-05*) |
| **Lysine** | 95 | 14 | 1.71 (0.2424) |
| **Methionine** | 134 | 44 | 56.39 (9.99e-05*) |
| **Glutamine** | 34 | 10 | 5.78 (0.0251) |
| **Tryptophan** | 62 | 9 | 1.07 (0.2936) |
| **TOTAL** | 387 | 105 | 15.96 (0.0072**) |

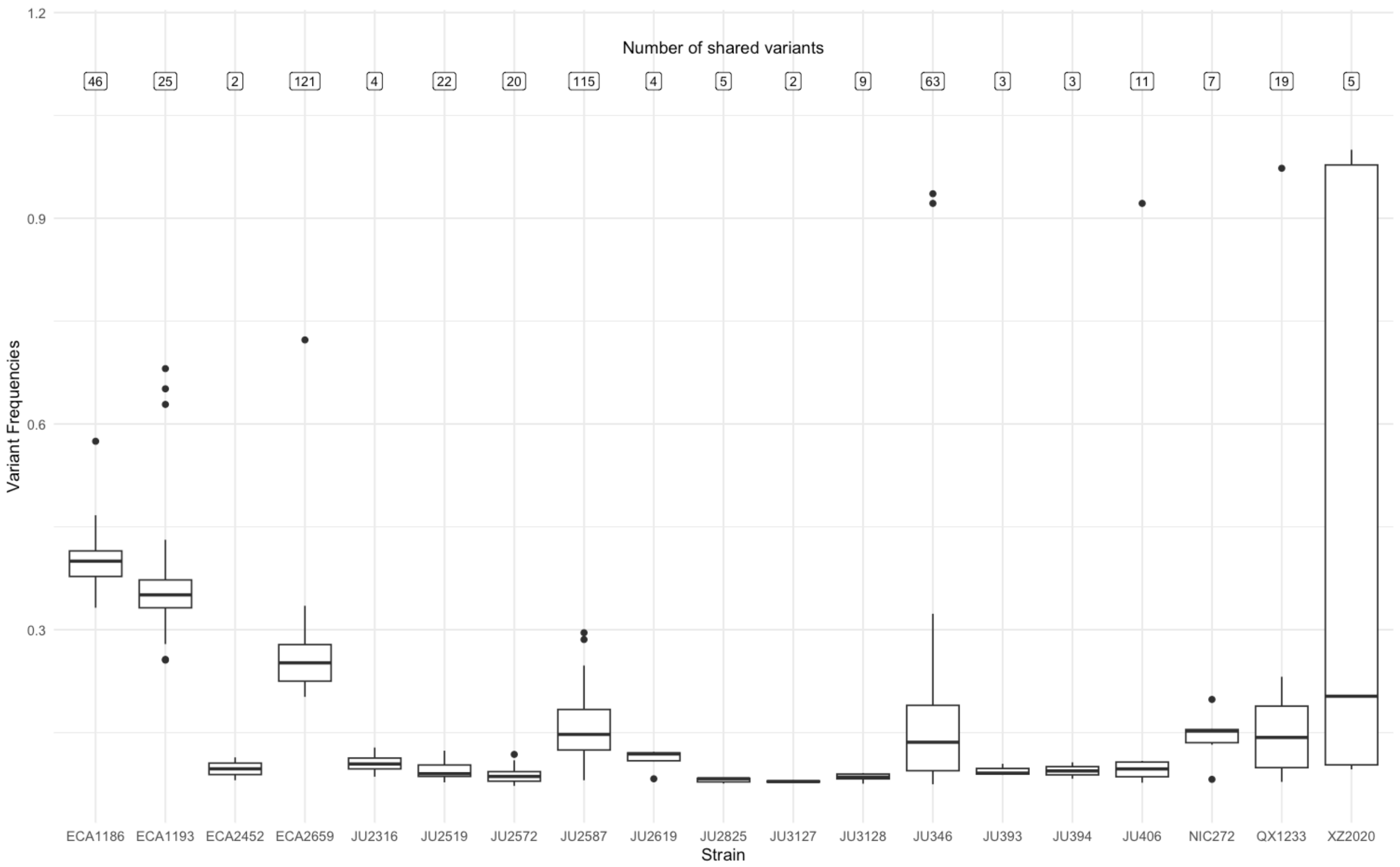


**Supplementary Fig. S1.** The frequencies of mtDNA variant calls in 19 *C. elegans* natural isolates from the CaeNDR database (Crombie et al. 2024) with >1 variant call per line in low frequency that were also shared with fixed variants in other natural isolates. The top of the figure lists the number of shared variants per line that are not fixed. We suspect that these calls did not result from true heteroplasmic variants but from DNA contamination from other natural isolates. Data from these lines was not included in the final analysis of the substitution spectrum in natural isolates.

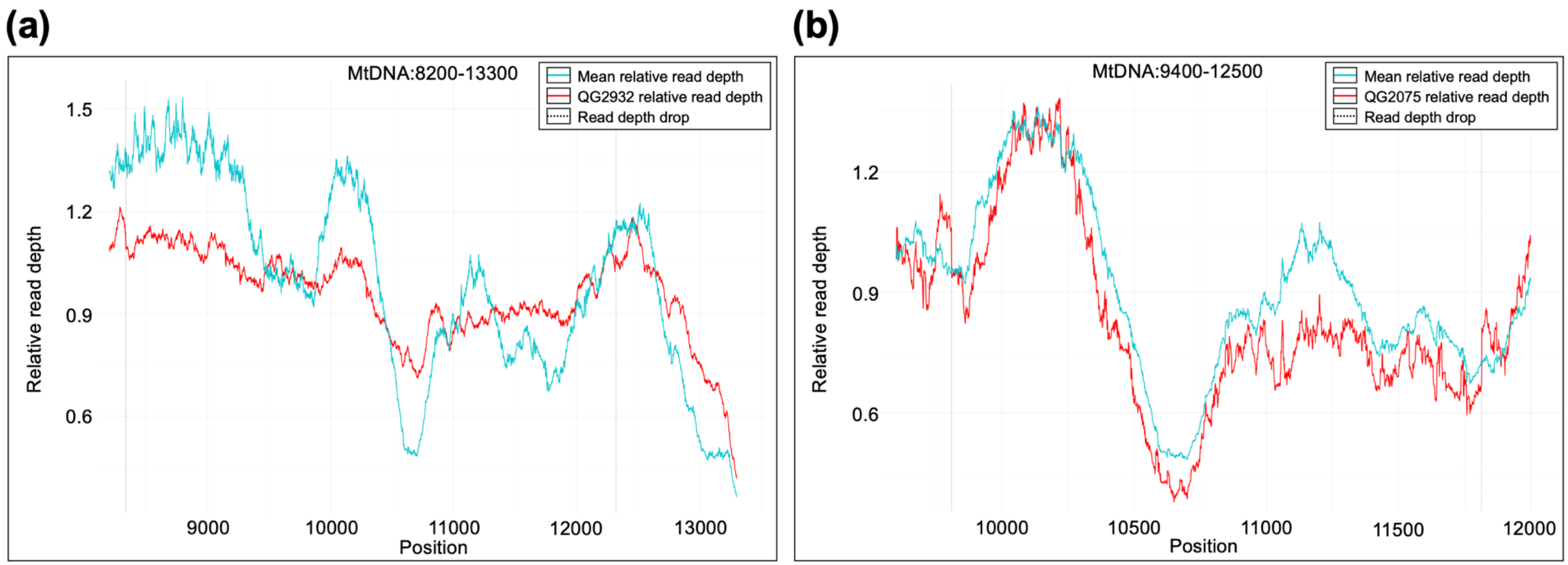


**Supplementary Fig. S2.** The distribution of read depth for two mitochondrial genomes with low heteroplasmic frequencies of mtDNA deletions. The horizontal axis represents the position in the reference strain mtDNA molecule and the vertical axis represents the relative read depth at any given position. The red and blue line is the distribution of relative read depth in the mtDNA harboring the deletion and mtDNA in all other natural isolates, respectively. The two grey vertical lines indicate the position of the deletion breakpoints which were identified independently in Delly, Manta and Pindel. a) Deletion in natural isolate QG2932. b) Deletion in natural isolate QG2075.

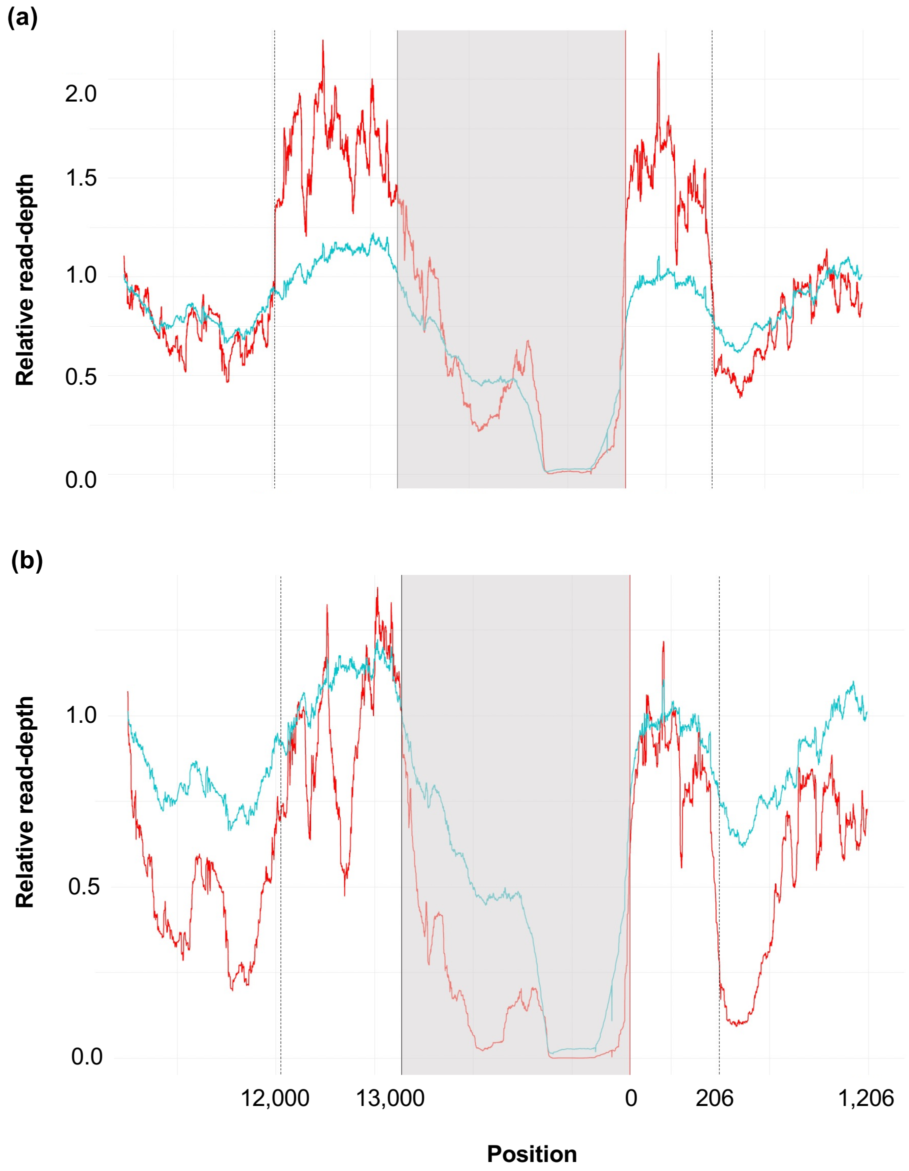


**Supplementary Fig. S3.** The distribution of read depth for two mitochondrial genomes with low heteroplasmic frequencies of mtDNA duplications. The breakpoints at the 5′ and 3′ ends of the duplications are shown in separate panels. The horizontal axis represents the position in the reference strain mtDNA molecule and the vertical axis represents the relative read depth at any given position. The red and blue lines represent the distribution of relative read depth in the mtDNA harboring the duplication and mtDNA in all other natural isolates, respectively. The two grey vertical lines indicate the position of the duplication breakpoints which were identified independently in Delly, Manta and Pindel. a) The left and right panels represent the 5′ and 3′ breakpoint of the duplication identified in natural isolate PS2025, respectively. b) The left and right panels represent the 5′ and 3′ breakpoint of the duplication identified in natural isolate MY25, respectively.
